# Hierarchical glycolytic pathways control the carbohydrate utilization regulator in human gut *Bacteroides*

**DOI:** 10.1101/2024.11.13.623061

**Authors:** Seth G. Kabonick, Kamalesh Verma, Jennifer L. Modesto, Victoria H. Pearce, Kailyn M. Winokur, Eduardo A. Groisman, Guy E. Townsend

## Abstract

Human dietary choices control the gut microbiome. Industrialized populations consume abundant amounts of glucose and fructose, resulting in microbe-dependent intestinal disorders. Simple sugars inhibit the carbohydrate utilization regulator (Cur), a transcription factor in members of the prominent gut bacterial phylum, *Bacteroidetes*. Cur controls products necessary for carbohydrate utilization, host immunomodulation, and intestinal colonization. Here, we demonstrate how simple sugars decrease Cur activity in the mammalian gut. Our findings in two *Bacteroides* species show that ATP-dependent fructose-1,6-bisphosphate (FBP) synthesis is necessary for glucose or fructose to inhibit Cur, but dispensable for growth because of an essential pyrophosphate (PPi)-dependent enzyme. Furthermore, we show that ATP-dependent FBP synthesis is required to regulate Cur in the gut but does not contribute to fitness when *cur* is absent, indicating PPi is sufficient to drive glycolysis in these bacteria. Our findings reveal how sugar-rich diets inhibit Cur, thereby disrupting *Bacteroides* fitness and diminishing products that are beneficial to the host.

## Introduction

Humans can consume over 3 times the recommended amounts of glucose and fructose, monosaccharides abundant in ultra-processed foods and beverages containing high fructose corn syrup^1^. Overconsumption of these sugars supersedes the absorptive capacity of the small intestine to reach the distal gut, where they are consumed by resident microbes^2^. High-sugar diets alter gut microbial composition and gene transcription, thereby increasing microbiota-dependent disease susceptibility^3,4^; however, the microbial processes that mediate sugar-rich, diet-dependent changes in the gut are not understood. Therefore, it is necessary to investigate the mechanisms by which refined sugar consumption reduces commensal fitness and disrupts beneficial microbiota-host interactions.

Carbon catabolite repression (CCR) is a global regulatory mechanism that prioritizes the utilization of preferred carbon sources over other available substrates^5-7^. In *Escherichia coli* (*Ec*), glucose and fructose impose CCR effects via the phosphoenolpyruvate (PEP):carbohydrate phosphotransferase system (PTS), which utilizes PEP to concomitantly import and phosphorylate target monosaccharides^8,9^. *Ec* employs the PTS to couple intracellular metabolite pools with sugar internalization and entry into central metabolism^10-12^. Ultimately, substrate phosphorylation by the PTS reduces cyclic adenosine monophosphate (cAMP) production, the allosteric activator of cAMP Receptor Protein (CRP), thereby reducing the expression of CRP-dependent genes, which mediate the utilization of less-preferred carbon sources^8,13^. Similar to CRP in *Ec*, *Bacteroides* encodes Cur, a transcription factor that is required for growth on several carbohydrates^14-16^; however, *Bacteroides* species lack both endogenous cAMP and PTS orthologs, indicating that a distinct CCR-like mechanism controls Cur activity^17-19^. Moreover, *Bacteroides* recognizes carbohydrates and initiates transcription in the periplasm before entering central metabolism^14-16^, further obscuring how sugars reduce Cur activity.

Cur regulates over 400 genes in the model organism, *Bacteroides thetaiotaomicron (Bt)^12^,* including products necessary for glycan utilization^20^, intestinal colonization^10,21^, and beneficial host interactions^4^. For example, *cur* is required for the expression of *fucI-fucR,* an operon that is necessary for fucose utilization^10,11^. Fucose is a constituent monosaccharide that decorates host mucosal glycans mediating a trans-kingdom signaling axis in the gut^22^. In addition, the *cur*-dependent gene, *fusA2*, encodes an alternative translation elongation factor that facilitates GTP-independent protein synthesis, a process necessary for intra-intestinal fitness^23^. Cur is also responsible for the expression of many polysaccharide utilization loci (PUL) that are important for providing *Bt* access to host- and dietary-derived nutrients. For example, PUL80 expression requires *cur* and contains *BT4295,* a putative chitobiase (*chb^PUL80^*) that directs immunotolerogenic T-cell development to reduce host-disease susceptibility^4^. Accordingly, a simple sugar diet reduces *chb^PUL80^* expression by inhibiting Cur activity, thereby disrupting T-cell differentiation and initiating colitis in mice^4^. Therefore, it is necessary to understand how simple sugars inhibit Cur activity for both bacterial fitness and host physiology.

Here, we report that Cur inhibition by glucose and fructose requires corresponding ATP-dependent hexokinases that facilitate their utilization. We establish that simple sugars impose CCR-like effects via a unique mechanism in *Bacteroides*, whereby phosphorylated sugars are converted into the ubiquitous metabolite – fructose-1,6-bisphosphate (FBP) – by distinct ATP- and PPi-dependent enzymes. Strikingly, ATP-dependent FBP synthesis is required for Cur inhibition by dietary sugars and necessary for *in vivo* fitness when *cur* is present, but otherwise dispensable for cell growth. In contrast, PPi-dependent FBP synthesis is essential for growth, indicating that PPi, rather than ATP, drives glycolysis in these organisms. Our findings identify unique pathways in *Bacteroides* that coordinate CCR-like effects on Cur activity in the presence of dietary sugars, thereby silencing colonization factor expression, hindering intra-intestinal fitness, and reducing products beneficial to host health.

## Results

### Glucose and fructose inhibit Cur activity in a dominant, dose-dependent manner

We previously demonstrated that *fusA2* is the most highly upregulated *cur*-dependent gene identified to date and Cur binding to the *fusA2* promoter region is necessary for *Bt* fitness in the murine gut^12^. To kinetically examine Cur activity during growth, we generated P-*fusA2* by introducing the *fusA2* promoter into *pBolux*, a reporter plasmid containing a *Bacteroides*-optimized luciferase cassette^24^. A *wild-type Bt* strain harboring P-*fusA2* exhibited higher bioluminescence than an isogenic strain harboring the promoter-less *pBolux* control plasmid in the *cur*-dependent substrates fucose (Fig. 1a) and N-acetylgalactosamine (galNAc) (Fig. 1b)^25^. In contrast, this strain exhibited decreasing bioluminescence during growth on the *cur*-independent monosaccharides, fructose or glucose (Fig. 1c & Extended Data Fig. 1a). When cultured in porcine mucosal O-glycans (PMOG), a mixture that increases *cur*-dependent gene transcription^10,20^, the *wild-type* strain harboring P-*fusA2* produced a 45-fold increase in bioluminescence by 18 hours (Fig. 1d). Bioluminescence from P-*fusA2* requires Cur activity because a *wild-type Bt* strain harboring P-*fusA2* but lacking the 22 base pair Cur binding site^12^ (P-*Δ22bp*) exhibited indistinguishable bioluminescence relative to strains harboring *pBolux* (Fig. 1a, b, c & Extended Data Fig. 1a, b, c). Similarly, a *cur-*deficient strain (*Δcur*) harboring P-*fusA2* produced bioluminescence comparable to *pBolux* when grown in identical conditions (Fig. 1c & Extended Data Fig. 1a, d, e). Bioluminescence from the *wild*-*type* strain harboring P-*fusA2* supplied PMOG and monosaccharide mixtures were altered according to the sugar identity. For example, mixtures containing galNAc initially reduced, but ultimately increased bioluminescence, whereas mixtures containing fucose initially had no effect but eventually increased bioluminescence above cells grown in PMOG alone (Extended Data Fig. 1f). Alternatively, a *wild-type* strain harboring P-*fusA2* supplied PMOG with increasing concentrations of fructose or glucose elicited corresponding reductions in bioluminescence (Fig. 1d & Extended Data Fig. 1g), similar to previously reported *cur*-dependent transcript levels^10^. Bioluminescence specifically reports changes in Cur activity because neither *Δcur* harboring P-*fusA2* nor the *wild-type* strain harboring P-*Δ22bp* produced increases compared to isogenic strains harboring p*Bolux* in all examined conditions (Extended Data Fig. 1b, c, d, e). Bioluminescence did not reduce cellular growth because strains harboring P-*fusA2* or P-*Δ22bp* grew comparably to isogenic strains harboring p*Bolux* in *cur*-dependent or -independent conditions (Extended Data Fig. 1h, i). Importantly, *cur* is dispensable for reporter functionality because a strain harboring P-*PUL22*^24^, a reporter containing the promoter region of the fructan utilization PUL (Extended Data Fig. 2a), exhibited bioluminescence resembling a *wild-type Bt* strain (Extended Data Fig. 2b) when cultured in PMOG supplemented with fructose, although slightly delayed due to a growth defect (Extended Data Fig. 2c). Collectively, these data establish that glucose and fructose inhibit Cur activity in a dominant and dose-dependent manner and that P-*fusA2* faithfully reports previously observed *cur*-dependent transcript levels^10^.

**Fig. 1.**
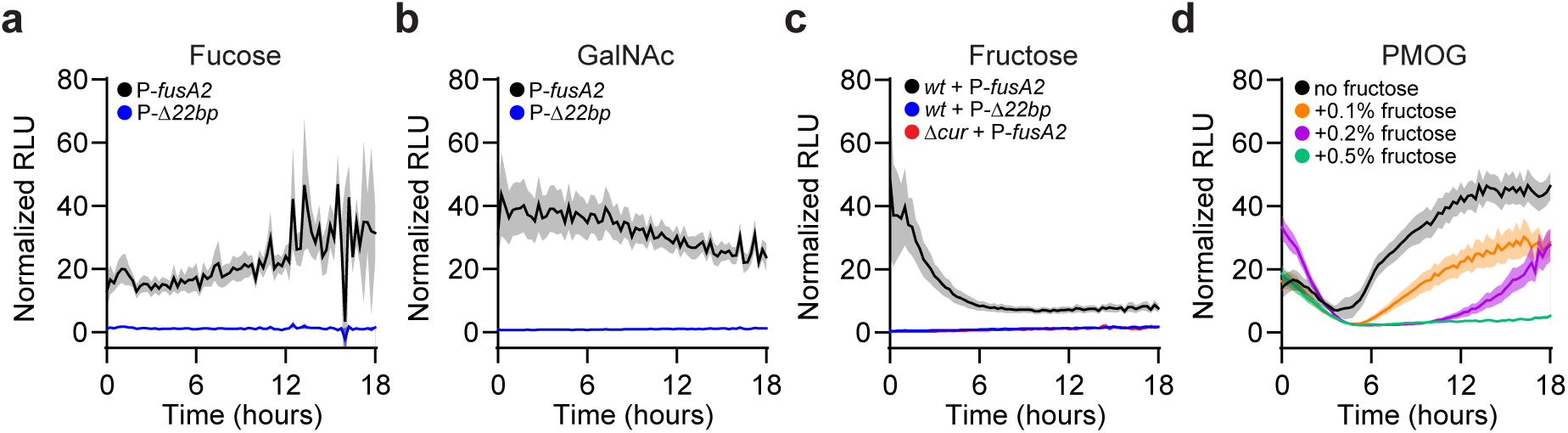
A bioluminescent transcriptional reporter of Cur activity. **a,b,** Bioluminescence from a *wild-type Bt* strain harboring P-*fusA2* (black) or P-*Δ22bp* (blue) cultured in media containing **(a)** fucose or **(b)** galNAc as the sole carbon source normalized to measurements collected from isogenic strains harboring a promoter-less p*Bolux* plasmid. **c,** Normalized bioluminescence from a *wild-type* strain harboring P-*fusA2* (black) or P-*Δ22bp* (blue) and a *Δcur Bt* strain harboring P-*fusA2* (red) cultured in fructose as a sole carbon source. **d,** Normalized bioluminescence from a *wild-type Bt* strain harboring P-*fusA2* cultured in PMOG as the sole carbon source (black) or in combination with 0.1% (orange), 0.2% (purple), or 0.5% (green) fructose. For panels **a-d,** n=8, error is SEM in color matched shading.

### Cur inhibition by fructose and glucose requires ATP-dependent substrate phosphorylation

Many other bacterial taxa alter transcription following sugar transport into the cytoplasm^26^; however, *Bacteroides* senses sugars prior to transport^14,24^, complicating the way fructose or glucose reduces Cur activity. For example, periplasmic fructose binds the sensor protein, BT1754 (Sensor^PUL22^), which is embedded in the inner membrane. Fructose binding to the sensor^PUL22^ directly controls the expression of PUL genes responsible for fructose transport via BT1763 (SusC^PUL22^) and putative phosphorylation by BT1757 (Frk^PUL22^) (Extended Data Fig. 2a)^14^. Consistent with these studies, a *wild-type Bt* strain harboring P-*PUL22* exhibited increased bioluminescence during growth in PMOG containing 0.2% fructose (Extended Data Fig. 3a) and the addition of fructose increased *susC^PUL22^* transcripts 607-fold (Extended Data Fig. 3b). In contrast, a *sensor^PUL22^*-deficient strain (*Δsensor^PUL22^*) produced no bioluminescence increases when harboring P-*PUL22* (Extended Data Fig. 3a) and *susC^PUL22^* transcript amounts did not significantly increase (Extended Data Fig. 3b), indicating that the Sensor^PUL22^ is required to transcribe genes necessary for fructose utilization, in agreement with previous findings^14,24^.

The Sensor^PUL22^ is also required for fructose-dependent Cur inhibition because *Δsensor^PUL22^* harboring P-*fusA2* produced similar bioluminescence during growth in PMOG alone or in combination with 0.5% fructose (Fig. 2a). Conversely, bioluminescence from *Δsensor^PUL22^* was reduced when grown in a mixture of PMOG and glucose (Fig. 2a), similar to *wild-type Bt* (Extended Data Fig. 1e). Furthermore, fructose decreased the abundance of *fusA2* and *chb^PUL80^* transcripts 181- and 747-fold, respectively, in the *wild-type* strain as previously reported^10^, but had no effect in the *Δsensor^PUL22^*mutant (Fig. 2b & Extended Data Fig. 3c). In contrast, glucose addition to PMOG-grown *Δsensor^PUL22^* reduced *fusA2* and *chb^PUL80^* transcripts 809-fold and 1,784-fold, respectively, similar to the 908-fold and 2,058-fold reductions exhibited by *wild-type Bt* under identical conditions (Fig. 2b & Extended Data Fig. 3c). Therefore, the Sensor^PUL22^ is required for Cur inhibition by fructose but not glucose.

**Fig. 2.**
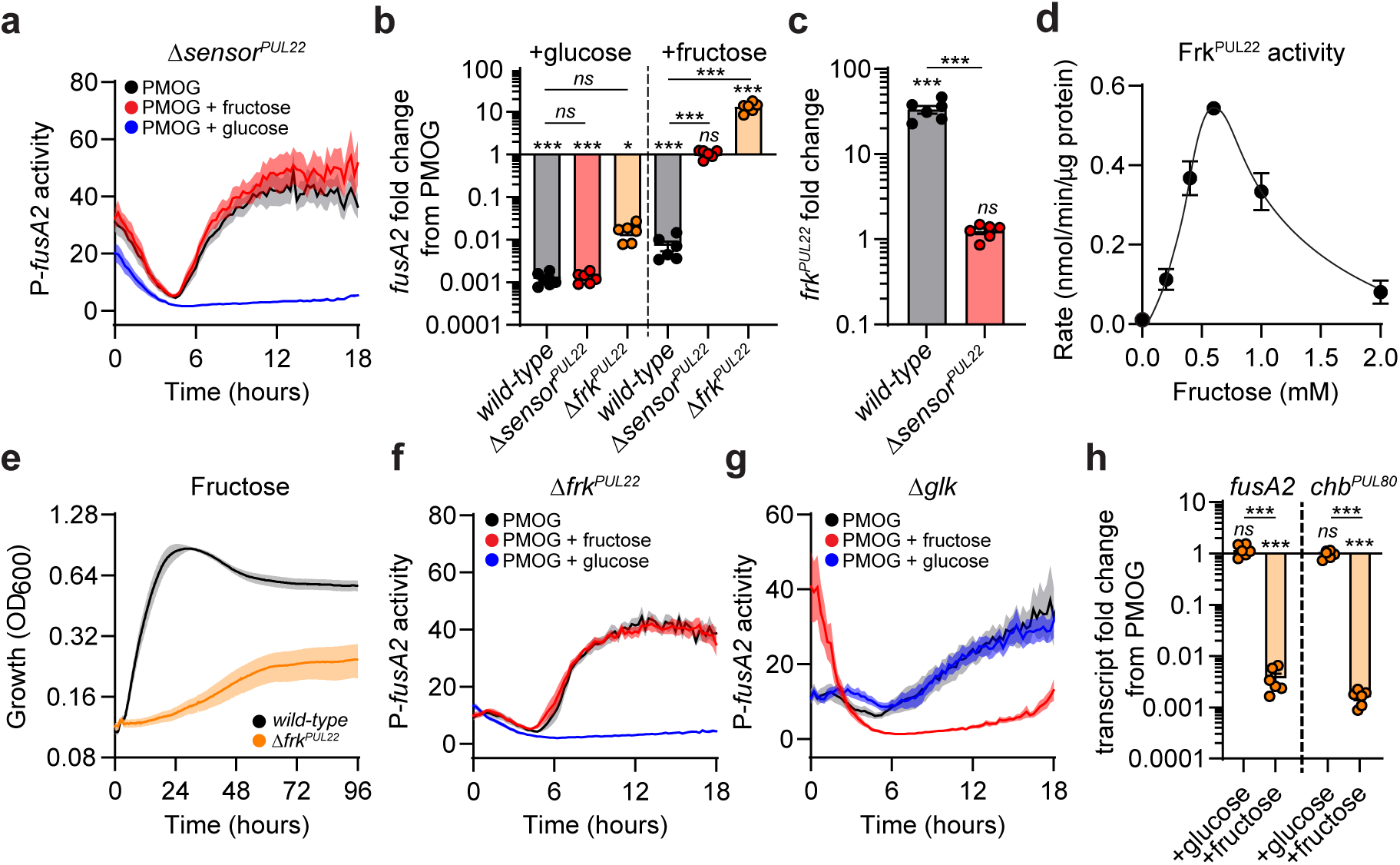
Fructose and glucose require phosphorylation for Cur inhibition. **a,** Normalized bioluminescence from *Δsensor-PUL22* harboring P-*fusA2* cultured in media containing PMOG (black), PMOG and fructose (red), or PMOG and glucose (blue). **b,** Fold change of *fusA2* transcript amounts from *wild-type*, *ΔsensorPUL22*, and *ΔfrkPUL22* cultured in media containing PMOG as the sole carbon source 10 minutes following the addition of 0.2% glucose (left) or 60 minutes after 0.2% fructose addition (right). **c,** Fold change of *frkPUL22* transcripts from *wild-type* or *ΔsensorPUL22* 60 minutes following the addition of 0.2% fructose to cells cultured in media containing PMOG. **d,** Fructokinase activity of purified FrkPUL22 protein. **e,** Growth of *wild-type* or *Δ frkPUL22* in minimal media containing fructose as the sole carbon source. **f,g,** Normalized bioluminescence from **(f)** *ΔfrkPUL22* or **(g)** *Δglk* harboring P-*fusA2* cultured in media containing PMOG (black) as sole carbon source or equal mixtures of PMOG and fructose (red) or glucose (blue). **h,** Fold change of *fusA2* (left) and *chbPUL80* (right) transcript amounts from *Δglk* cultured in media containing PMOG as the sole carbon source 10 minutes following 0.2% glucose addition or 60 minutes following fructose addition. For panels **a,e-g,** n=8; error is SEM in color matched shading. For panels **b,c,h,** n=6; error is SEM. For panel **d,** n=4; error is SEM. P-values were calculated by 2-way ANOVA with Fisher’s LSD test and * represents values < 0.05, ** < 0.01, *** < 0.001.

The PUL22 regulon includes *frk^PUL22^*, which is putatively responsible for the phosphorylation of fructose as it enters the cytoplasm^25^. Accordingly, *frk^PUL22^* transcripts increased 33-fold in a *sensor^PUL22^*-dependent manner following the introduction of fructose (Fig. 2c), Frk^PUL22^ protein possessed *in vitro* fructokinase activity (Fig. 2d), and a *frk^PUL22^*-deficient strain (*Δfrk^PUL22^*) was unable to grow on fructose (Fig. 2e), indicating that this enzyme is likely the sole *Bt* fructokinase. In contrast, *Δfrk^PUL22^* did not exhibit a growth defect in glucose (Extended Data Fig. 3d) or PMOG (Extended Data Fig. 3e). Fructose phosphorylation is required for entry into central metabolism, but not necessary for Sensor^PUL22^-directed transcription, because *Δfrk^PUL22^* harboring P-*PUL22* exhibited increased bioluminescence (Extended Data Fig. 3a) and *susC^PUL22^*transcript amounts increased 210-fold in *Δfrk^PUL22^* following the introduction of fructose (Extended Data Fig. 3b). Conversely, *susC^PUL22^*transcript amounts did not increase in *Δsensor^PUL22^* (Extended Data Fig. 3b). Frk^PUL22^ is necessary for fructose-dependent Cur inhibition because *Δfrk^PUL22^* harboring P-*fusA2* produced similar bioluminescence during growth in PMOG or a combination of PMOG and fructose but exhibited reduced bioluminescence during growth in PMOG and glucose (Fig. 2f), similar to *wild-type Bt* (Extended Data Fig. 1d) and *Δsensor^PUL22^* (Fig. 2a). Thus, fructose specifically decreases Cur activity via this compartmentalized signaling machinery by increasing fructokinase transcription, suggesting that sugar phosphorylation is necessary to reduce Cur activity.

We predicted that glucose phosphorylation mediates Cur inhibition by a distinct mechanism, likely involving a hexokinase, because glucose addition reduced Cur activity independent of Frk^PUL22^ (Fig. 2b and Fig. 2f). BT2493 (Glk) exhibits 92% identity to RokA from *Bacteroides fragilis* (*Bf*), which is necessary for *Bf* growth on glucose^27^. We determined that Glk exhibits glucokinase activity *in vitro* (Extended Data Fig. 4a) and the growth of a *glk*-deficient strain (*Δglk*) was severely impaired on glucose as the sole carbon source but grew similarly to *wild-type Bt* in media containing fructose or PMOG (Extended Data Fig. 4b). Accordingly, *Δglk* harboring P-*fusA2* exhibited similar bioluminescence during growth in media containing PMOG or equal amounts of PMOG and glucose (Fig. 2g). Neither *fusA2* nor *chb^PUL80^*transcripts decreased following the addition of 0.2% glucose to *Δglk* growing on PMOG (Fig. 2h); however, fructose-dependent silencing resembled *wild-type Bt* (Fig. 2b), indicating that Glk is necessary for glucose, but not fructose, to inhibit Cur activity. Therefore, glucose and fructose require distinct ATP-dependent hexokinases for their utilization and inhibition of Cur activity.

### *Bt* possesses both ATP- and PPi-dependent glycolytic pathways

In glycolysis, phosphoglucoisomerase (Pgi) converts glucose-6-phosphate (G6P) into fructose-6-phosphate (F6P), which is subsequently phosphorylated into FBP (Fig. 3a). Because FBP synthesis is the primary regulatory step in glycolysis and *wild-type Bt* cells exhibit reduced FBP amounts^28^ with corresponding increases in Cur activity during carbon limitation^10,12^, we hypothesized that glucose and fructose reduce Cur activity via this glycolytic step. To test this, we first examined the biochemical activities of 3 putative *Bt* phosphofructokinase (Pfk) enzymes: BT2062 (PfkA), BT1102 (PfkB), and BT3356 (PfkC), whose amino acid sequences share greater than 39% identity with *Ec* PfkA (Extended Data Fig. 5a). We determined that PfkA and PfkB, but not PfkC, are *bona fide* ATP-dependent Pfks, because expression of each in a *pfk*-deficient *Ec* strain^29^ restored growth on glucose (Extended Data Fig. 5b) when expressed in an inducer-dependent manner (Extended Data Fig. 5c). Moreover, purified recombinant PfkA and PfkB proteins exhibited activity *in vitro*, whereas PfkC did not (Fig. 3b). Therefore, like *Ec, Bt* encodes two Pfk enzymes that convert F6P into FBP using ATP (Fig. 3a).

**Fig. 3.**
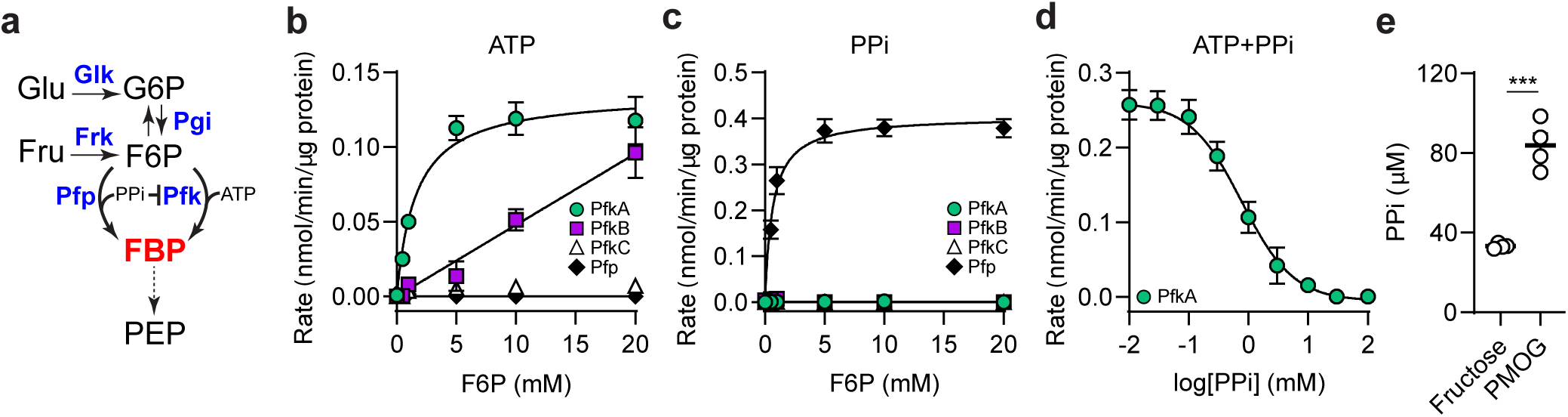
Distinct enzyme classes synthesize FBP in *Bt*. **a,** Schematic of the glycolytic pathway in *Bt*. **b,c,** Enzyme kinetics of purified PfkA (green circles), PfkB (purple squares), PfkC (white triangles), or Pfp (black diamonds) in reactions containing either **(b)** ATP or **(c)** PPi as a phosphoryl donor. **d,** PfkA activity in the presence of increasing PPi amounts. **e,** PPi amounts in whole cell lysates of *wild-type* grown in fructose or PMOG as the sole carbon source. For panel **a**, abbreviations are as follows: glucose (Glu), fructose (Fru), glucose-6P (G6P), fructose-6P (F6P), fructose bisphosphate (FBP), phosphoenolpyruvate (PEP), adenosine triphosphate (ATP), pyrophosphate (PPi), phosphofructokinase (Pfk), phosphofructose phosphotransferase (Pfp), phosphoglucose isomerase (Pgi), glucokinase (Glk), fructokinase (Frk). For panels **b-d,** n=4, error is SEM. For panel **e,** n=4, error is SEM; *P*-values were calculated by 1-way ANOVA with Fisher’s LSD test and *** represents values < 0.001.

*Bacteroides* species also encode a putative fructose-6-phosphate 1-phosphotransferase, BT0307 (Pfp), which synthesizes FBP independent of ATP by utilizing pyrophosphate (PPi) as a phosphoryl donor^30^. BT0307 exhibited PPi-dependent FBP synthesis *in vitro*, whereas PfkA and PfkB did not (Fig. 3c). *Bt* expresses *pfkA*, *pfkB*, and *pfp* simultaneously during growth in PMOG and the addition of glucose or fructose decreased *pfkB* and increased *pfp* 2-fold, respectively, but had no effect on *pfkA* transcript amounts (Extended Data Fig. 5d). These results suggest that both ATP- and PPi-dependent FBP synthesis occur simultaneously in the cell; however, PfkA activity was potently inhibited by PPi *in vitro* (Fig. 3d), suggesting its activity is governed by intracellular PPi amounts. Furthermore, PMOG grown *wild-type Bt* exhibited 2.5-fold higher PPi amounts than in fructose-containing media (Fig. 3e), suggesting that fructose utilization permits ATP-dependent FBP synthesis by reducing PPi levels below inhibitory concentrations. Thus, in contrast to other enteric bacteria, *Bt* employs divergent FBP biosynthetic pathways.

### ATP-dependent FBP synthesis controls Cur activity

To investigate the role of FBP synthesis in Cur inhibition, we constructed strains lacking *pfkA* (*ΔpfkA*), *pfkB* (*ΔpfkB*), or both *pfkA* and *pfkB* genes (*ΔpfkAB*). *ΔpfkA and ΔpfkB* grew similarly to *wild-type Bt* in fructose or PMOG as sole carbon sources (Fig. 4a, Extended Data Fig. 6a), whereas *ΔpfkA* had slightly increased growth rates and maxima in glucose (Extended Data Fig. 6b, c). Unexpectedly, *ΔpfkAB* grew similarly to *wild-type Bt* on fructose (Fig. 4a), glucose, or PMOG (Extended Data Fig. 6a, b, c), implying that ATP-dependent FBP synthesis is dispensable for *in vitro* growth. Cell extracts prepared from *ΔpfkA* or *ΔpfkAB* exhibited no Pfk activity, indicating that ATP-dependent FBP synthesis was entirely absent in these strains (Fig. 4b). In contrast, extracts prepared from *wild-type* and *ΔpfkA* produced indistinguishable PPi-dependent FBP synthesis *in vitro* at rates 11.2- and 38.5-fold greater than ATP-dependent reactions, respectively (Fig. 4b). Conversely, we were unable to generate a strain lacking the *BT0307* open-reading frame, suggesting *pfp* is necessary for growth (Extended Data Fig. 6d), which agrees with transposon-based examinations of essential genes in *Bt*^31,32^. Therefore, PPi-dependent FBP synthesis provides sufficient metabolic flux, thereby rendering ATP-dependent FBP synthesis dispensable for growth.

**Fig. 4.**
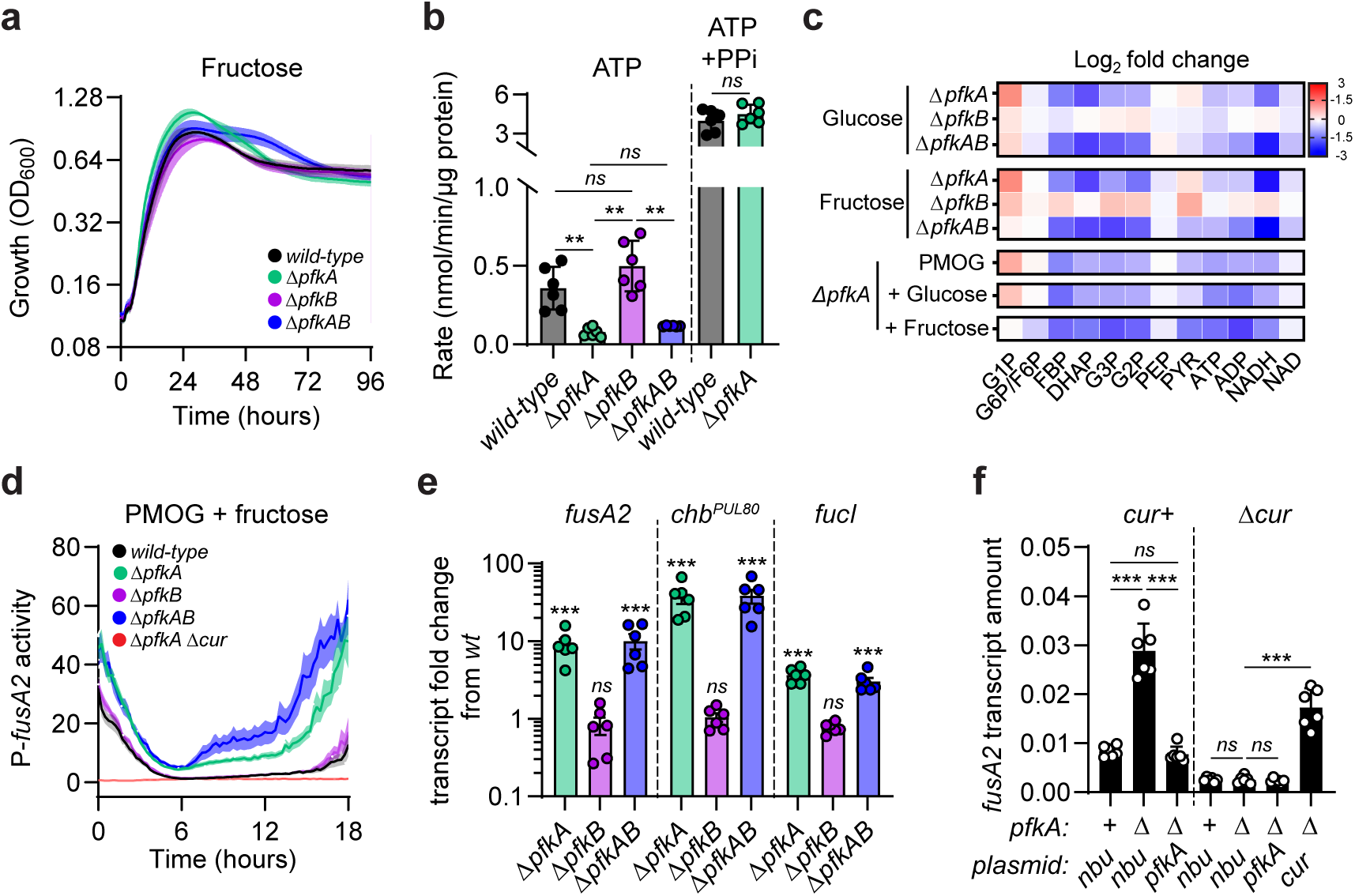
ATP-dependent FBP production is required for Cur inhibition but dispensable for growth. **a,** Growth of *wild-type* (black)*, ΔpfkA* (green), *ΔpfkB* (purple), or *ΔpfkAB* (blue) in minimal media containing fructose as a sole carbon source. **b,** FBP synthesis measured from whole cell lysates of *wild-type, ΔpfkA*, *ΔpfkB*, or *ΔpfkAB*. **c,** Log fold change of steady-state glycolytic intermediates in *ΔpfkA*, *ΔpfkB*, or *ΔpfkAB* in glucose, fructose, PMOG, PMOG2 with glucose, or PMOG with fructose. **d,** Bioluminescence from *wild-type* (black), *ΔpfkA* (green), *ΔpfkB* (purple), *ΔpfkAB* (blue), or *ΔpfkA Δcur* (red) harboring P-*fusA2* in minimal media containing equal amounts of PMOG and fructose. **e,** Fold change of *fusA2, chbPUL80, and fucI* transcript amounts relative to *wild-type* from *ΔpfkA*, *ΔpfkB*, or *ΔpfkAB* cultured in minimal media containing fructose as a sole carbon source. **f,** *fusA2* transcript amounts in *wild-type*, *ΔpfkA*, *Δcur*, and *ΔpfkA Δcur* harboring empty vector (*nbu*) or complementing plasmids grown in glucose as the sole carbon source. For panel **c,** abbreviations are as follows: glucose-1P (G1P), glucose-6P (G6P), fructose bisphosphate (FBP), dihydroxyacetone-P (DHAP), 3-phosphoglycerate (G3P), 2-phosphoglycerate (G2P), phosphoenolpyruvate (PEP), pyruvate (PYR), adenosine triphosphate (ATP), adenosine diphosphate (ADP), nicotinamide adenine dinucle-otide (NAD). For panels **a,d,** n=8; error is SEM in color matched shading. For panel **b,** n=4; error is SEM. For panels **e,f,** n=6; error is SEM. For panels **b,e,f,** *P*-values were calculated by 2-way ANOVA with Fisher’s LSD test and * represents values < 0.05, ** < 0.01, *** < 0.001.

To determine how Pfk enzymes support glycolysis, we measured steady-state metabolite abundances in *wild-type*, *ΔpfkA, ΔpfkB,* and *ΔpfkAB Bt* strains. Accordingly, *ΔpfkA* and *ΔpfkAB* exhibited respective 2.9- and 2.7-fold lower steady-state FBP amounts in *Bt* grown on glucose and fructose, respectively, whereas *ΔpfkB* showed no significant changes in either condition (Fig. 4c & Extended Data Fig. 7a) indicating that PfkA is the dominant ATP-dependent FBP biosynthetic enzyme in *Bt*. Downstream glycolytic metabolites, such as dihydroxyacetone phosphate (DHAP), also exhibited corresponding reductions in *ΔpfkA* but not *ΔpfkB* grown in either glucose or fructose (Fig. 4c & Extended Data Fig. 7b). Conversely, PEP and pyruvate amounts were similar across all strains and conditions (Fig. 4c & Extended Data Fig. 7c, d), suggesting these metabolites are maintained by processes independent of PfkA and PfkB. Additionally, *ΔpfkAB* resembled *ΔpfkA* across all metabolites and conditions (Fig. 4c & Extended Data Fig. 7a-d). Finally, ATP-dependent FBP synthesis is necessary for maintaining steady-state nucleotide triphosphates (NTP) and reducing equivalents because ATP and NADH amounts were also reduced in *ΔpfkA* and *ΔpfkAB* (Fig. 4c).

To determine if ATP-dependent FBP synthesis contributes to Cur inhibition, we measured bioluminescence in isogenic *wild-type*, *ΔpfkA*, *ΔpfkB*, and *ΔpfkAB* harboring P-*fusA2*. Strikingly, *ΔpfkA* exhibited a 9.2-fold increase in bioluminescence by 12 hours during growth in equal amounts of fructose and PMOG (Fig. 4d) and increased 10.5-fold by the same time point in glucose and PMOG (Extended Data Fig. 8a). Bioluminescence increased further in *ΔpfkAB* over *ΔpfkA,* however, *ΔpfkB* exhibited bioluminescence comparable to *wild-type Bt* harboring an identical reporter plasmid during growth on either mixture (Fig. 4d & Extended Data Fig. 8a). Similarly, *ΔpfkA* and *ΔpfkAB* produced greater bioluminescence than *wild-type* during growth on glucose, fructose, and PMOG as sole carbon sources (Extended Data Fig. 8b, c, d), suggesting that disabling ATP-dependent FBP synthesis increases Cur activity.

Consistent with observed bioluminescence increases, *pfkA*, but not *pfkB*, is required for Cur inhibition by glucose and fructose because the *cur-*dependent transcripts *fusA2*, *chb^PUL80^*, and *fucI* increased 9.1-, 37.2-, and 3.7-fold respectively in *ΔpfkA* when grown on fructose as the sole carbon source, whereas *ΔpfkB* exhibited transcript amounts indistinguishable from *wild-type* (Fig. 4e). Similarly, *ΔpfkA* exhibited respective 2.7-, 6-, and 1.7-fold increased *fusA2* (Fig. 4f), *chb^PUL80^* (Extended Data Fig. 8e), and *fucI* (Extended Data Fig. 8f) transcript amounts during growth in glucose that were abolished when the *pfkA* gene was supplied *in trans* (Fig. 4f, Extended Data Fig. 8e, f). The amounts of all three transcripts were indistinguishable between *Δcur* and *ΔpfkA Δcur*; but increased in *ΔpfkA Δcur* complemented *in trans* with *cur* but not *pfkA* (Fig. 4f, Extended Data Fig. 8e, f).

To explore the possibility of glycolytic intermediates directly impacting Cur’s ability to bind its regulated promoters, we performed electromobility shift assays (EMSA) with purified Cur protein and the *wild-type fusA2* promoter or a *Δ22*bp control probe. Increasing amounts of Cur protein shifted the *wild-type fusA2* probe in a dose-dependent manner, with migrations reflecting putative di-, tetra-, and oligo-meric interactions. Conversely, increasing Cur amounts did not alter the migration of the *Δ22*bp probe (Extended Data Fig. 9a-b), indicating that shifting observed with the *wild-type* probe was specific. Glycolytic intermediates that were differentially abundant in *ΔpfkA* are likely indirectly responsible for altering Cur activity because *fusA2* probe shifting was unaltered when F6P, FBP, DHAP, and PEP were included in the assay (Extended Data Fig. 9c). Co-incubation with *Bt* cell lysate fractions also did not change probe migration, suggesting that ligand-dependent probe binding may require different concentrations, additional co-factors, or distinct *in vitro* conditions (Extended Data Fig. 9d).

### ATP-dependent FBP synthesis regulates Cur *in vivo*

We hypothesized that PfkA may play a role in intra-intestinal *Bt* fitness because *ΔpfkA* exhibited increased *cur*-dependent products in the presence of simple sugars. Therefore, we inoculated germ-free mice with equal amounts of *wild-type* and *ΔpfkA* and followed their abundance in fecal contents over time*. pfkA* confers a fitness advantage *in vivo* because the relative abundance of *ΔpfkA* decreased 135-fold after 2 weeks (Fig. 5a). This suggests that the demands of intra-intestinal life require ATP-dependent FBP production even though this process reduces growth rates and maxima *in vitro* (Fig. 4a & Extended Data Fig. 6b, c). We also examined the relative abundances of *Δcur* and *ΔpfkA Δcur* following co-introduction into germ-free mice. Strikingly, both strains were present at nearly indistinguishable abundances across two weeks, suggesting that the sole purpose of *pfkA* is to regulate Cur in the mammalian intestine (Fig. 5b).

**Fig. 5.**
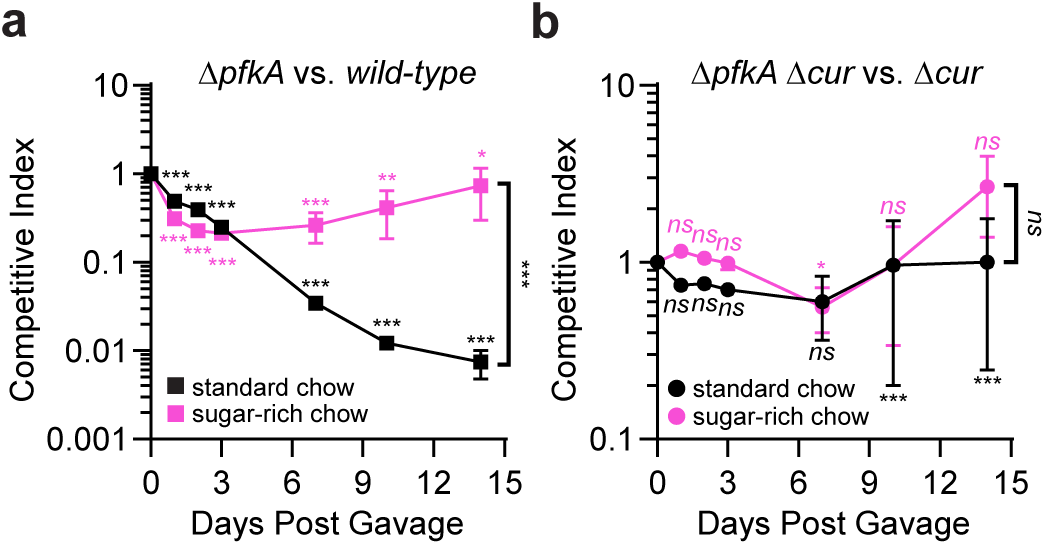
ATP-dependent FBP production is required for intestinal fitness by controlling Cur. **a,** Competitive fitness of *ΔpfkA* co-introduced into germ-free mice with equal amounts of *wild-type* and fed a standard polysaccharide rich chow (black squares, n=10) or a sugar-rich chow (pink squares, n = 8). **b,** Competitive fitness of *ΔpfkA Δcur* co-introduced into germ-free mice with equal amounts of *Δcur* and fed a standard polysaccharide rich chow (black circles, n=10) or a sugar-rich chow (pink circles, n=4). P-values were calculated using 2-way ANOVA with Bonferroni correction and * indicates values < 0.05, ** < 0.01, *** < 0.001.

Because PfkA is required for reducing *cur-*dependent products (Fig. 4f and Extended Data Fig. 8e, f), we reasoned that a high sugar diet, primarily comprised of glucose and sucrose, would ameliorate the *ΔpfkA* fitness defect by inhibiting Cur activity in *wild-type Bt*. Accordingly, *ΔpfkA* abundance was reduced only 1.4-fold compared to *wild-type Bt* after 2 weeks in mice provided a sugar-rich chow (Fig. 5a). This indicates that dietary sugar inhibits Cur activity *in vivo* and reduces the *wild-type* strain’s advantage over *ΔpfkA*. Finally, sugar-dependent effects on Cur activity are responsible for decreased *Bt* fitness because the abundances of *Δcur* and *ΔpfkA Δcur* were indistinguishable following introduction into germ-free mice fed a sugar-rich diet (Fig. 5b). Taken together, our data demonstrates that ATP-dependent FBP synthesis benefits *Bt* by regulating Cur activity in the mammalian gut and that PfkA activity is required for dietary sugars to inhibit Cur.

### PfkA regulates Cur activity across human intestinal *Bacteroides* species

Along with *Bt*, human *Bacteroides* isolates *Bf*, *Bacteroides ovatus*, and *Phocaeicola vulgatus* employ Cur to regulate their corresponding *fusA2* orthologs and *Bf* exhibits identical transcript silencing in response to glucose or fructose addition during growth in PMOG^10^. Each species also encodes *pfkA* orthologs (*BF9343_3444*, *BACOVA_00639*, and *BVU_1935*, respectively) and *pfp* orthologs (*BF9343_2852*, *BACOVA_01648*, and *BVU_2286*, respectively) suggesting that ATP-dependent FBP synthesis could govern Cur activity across *Bacteroides* species. We introduced P*-Bf-fusA2 (*p*Bolux* containing the 300 bp region preceding *BF9343_3536)* into *wild-type Bf* and isogenic strains lacking the *pfkA*-ortholog, *BF9343_3444* (*ΔBf-pfkA*) or *cur*-ortholog, *BF9343_0915* (*ΔBf-cur*). A *wild-type* strain harboring P-*Bf-fusA2* exhibited a 12.3-fold increase in bioluminescence compared to a strain harboring a promoter-less p*Bolux* during growth in PMOG (Extended Data Fig. 10a), displaying trends similar to *wild-type Bt* harboring P-*fusA2* (Fig. 1d). In contrast, an identical strain exhibited 2.3-fold increased bioluminescence compared to the promoter-less control strain during growth in either glucose (Extended Data Fig. 10b) or fructose (Extended Data Fig. 10c) as the sole carbon source. The addition of glucose (Fig. 6a) or fructose (Fig. 6b) to PMOG reduced bioluminescence from *wild-type* harboring P-*Bf-fusA2,* whereas *ΔBf-pfkA* displayed increased bioluminescence under all conditions (Fig. 6a, b & Extended Data Fig. 10a, b, c). As expected, cell extracts from *ΔpfkA* exhibited no ATP-dependent FBP synthesis *in vitro* compared to *wild-type* (Fig. 6c), collectively indicating that this process inhibits Cur activity in *Bf*. Consistent with this notion, bioluminescence from a *Bf* strain lacking both *pfkA* and *cur* (*ΔBf-pfkA ΔBf-cur*) was indistinguishable from *ΔBf-cur*, which were lower than isogenic strains harboring the promoter-less p*Bolux* plasmid (Fig. 6a, b, and Extended Data Fig. 10a, b, c). Therefore, Cur inhibition by ATP-dependent FBP production during growth in dietary sugars is conserved among human *Bacteroides* species.

**Fig. 6.**
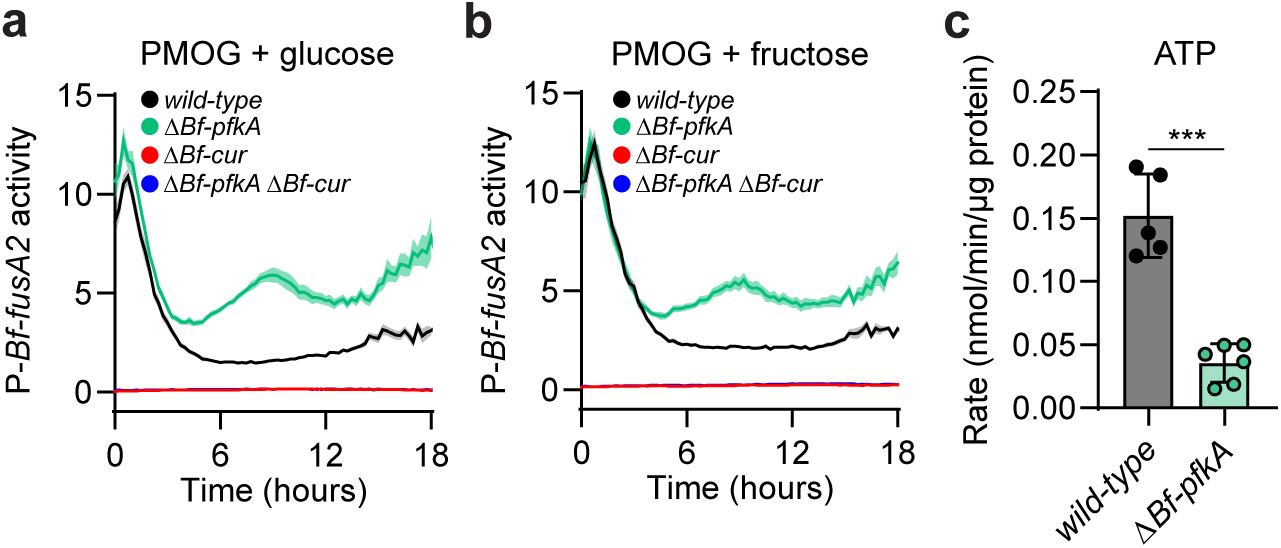
*Bf-pfkA* is required for glucose- and fructose-mediated Cur inhibition and ATP-dependent FBP synthesis in *B. fragilis*. **a,b,** Bioluminescence from *wild-type* (black)*, ΔBf-pfkA* (green), *ΔBf-cur* (red), or *ΔBf-pfkA ΔBf-cur* (blue) harboring P*-Bf-fusA2* cultured in media containing a mixture of equal amounts of PMOG and **(a)** glucose or **(b)** fructose. **c,** FBP synthetic rates measured from whole cell lysates of *wild-type* or *ΔBf-pfkA* supplied ATP as a phosphoryl donor. For panels **a,b,** n=8 and error is SEM in color matched shading. For panel **c,** n=5 and error is SEM. P-values were calculated using an unpaired t-test and *** represents values < 0.001.

## Discussion

Our work reveals a pathway in human intestinal *Bacteroides* species required for glucose and fructose to inhibit Cur. We determined that phosphorylation of either monosaccharide requires ATP-dependent sugar kinases necessary for growth, a strategy distinct from traditional PTS that utilizes PEP to phosphorylate sugars upon transport (Fig. 2). Furthermore, we show that *Bt* possesses both ATP- and PPi-dependent enzymes, Pfk and Pfp respectively, which form a hierarchical paradigm to regulate FBP synthesis (Fig. 3) and control Cur activity in response to nutrient availability. We demonstrate that eliminating Pfk alleviates sugar-dependent Cur inhibition without reducing growth (Fig. 4); however, *pfkA* is necessary to control Cur during intestinal colonization (Fig. 5). Furthermore, ATP-dependent FBP synthesis inhibits Cur *in vivo* because the competitive defect exhibited by *ΔpfkA* (Fig. 5b) is abolished when simple sugars are abundant in the host diet. Finally, we established that these processes are conserved in *Bf* and are likely shared among the *Bacteroides* (Fig. 6).

Collectively, these data reveal mechanisms that *Bacteroides* species employ to navigate the diverse buffet of intestinal glycans and dietary sugars present in the distal gut. Our findings indicate that Cur activity is tuned based on the type and amount of sugars accessible to *Bacteroides* (Fig. 1d, Extended Data Fig. 1f, g). Furthermore, Cur regulation is likely important to prioritize the metabolism of preferred sugars to compete with other gut bacteria for a limited supply of nutrients. Our findings explain how glucose and fructose consumption hinder *Bacteroides* intestinal fitness by exerting CCR-like effects on Cur activity via two hierarchical FBP biosynthetic pathways. Remarkably, PPi-dependent FBP synthesis occurs at a nearly 12-fold higher rate than ATP-dependent reactions; yet eliminating ATP-dependent synthesis when glucose or fructose is readily available reduced steady-state FBP levels by more than half (Extended Data Fig. 7a). To explain this phenomenon, we propose that the utilization of these simple sugars reduces PPi levels below an inhibitory threshold, thereby simultaneously diminishing the activity of Pfp and allowing for PfkA activity. Furthermore, the essentiality of *pfp*^33^ (Extended Data Fig. 6d) and lack of native pyrophosphorylase^30^ imply that *Bacteroides* have evolved to rely on PPi, rather than ATP, as a significant glycolytic energy source to conserve ATP. A similar energy-conservation strategy is employed in *Bt* via the *cur*-dependent gene, *fusA2*, whose product preserves NTP pools by facilitating GTP-independent protein synthesis^23^. A deeper examination into the role of PPi-dependent reactions across gut commensals is necessary to understand how bacteria balance NTP and PPi ratios to govern intestinal colonization.

FBP is a ubiquitous metabolite that serves as both a glycolytic intermediate and metabolic regulator of other crucial energy pathways^34^. Because endogenous FBP levels are inversely related to Cur activity^28^, but do not directly alter its binding to target promoters (Extended Data Fig. 9c), we hypothesize that FBP regulates other important energy pathways in *Bacteroides.* For example, FBP is required to stimulate glycogen production in the *Bacteroidetes* family member, *Prevotella bryantii*^35^. Intriguingly, *Bt* genes encoding putative glycogen metabolic enzymes are important fitness determinants in the murine intestine^31,32^ and are necessary for fitness in *cur*-dependent, but not -independent, substrates^33^. In light of this, we propose that host consumption of abundant glucose and fructose reconfigures central metabolism by stimulating ATP-dependent FBP synthesis and glycogenesis, in turn reducing an unknown Cur activator, and ultimately disrupting products necessary for *Bacteroides* fitness and beneficial host interactions.

## Methods

### Bacterial strains and growth conditions

All bacteria were cultured as described previously^10^ except for *Δglk* strains, which were cultured on rich and minimal media preparations where glucose was replaced with identical concentrations of fructose. Briefly, *Bacteroides* strains were cultured on brain-heart infusion agar (MilliporeSigma) containing 5% horse blood (Hardy) under anaerobic conditions (85% N_2_, 12.5% CO_2_, 2.5% H_2_). Liquid cultures were inoculated from a single colony into TYG media and incubated under anaerobic conditions. All bacterial strains included the following antibiotics where appropriate: 100 μg/mL ampicillin, 200 μg/mL gentamicin, 2 μg/mL tetracycline, or 25 μg/mL erythromycin (MilliporeSigma).

### Engineering chromosomal deletions

Indicated *Bt* genomic deletions were generated using pEXCHANGE-*tdk* plasmids harboring flanking sequences, amplified using the primers listed in Supplementary Table 2, as previously described^36^. In short, pEXCHANGE constructs were introduced via di-parental mating and chromosomal integration was validated by PCR. Parent strains were counter-selected on solid media containing 200ng/mL 5-fluoro-2-deoxyuridine (DOT Scientific). Indicated *Bf* genomic deletions were generated using pLGB13 plasmids harboring flanking sequences, amplified using the primers listed in Supplementary Table 2, as previously described^10,37^. All strains were validated using linear amplicon sequencing.

### Bacterial growth assays

Bacterial growth was measured as previously described^10^. Briefly, strains were cultured in anaerobic conditions to stationary phase in TYG and diluted 1:200 into reduced minimal media containing the carbon source of interest in 96- or 384-well clear microplates (Corning). Cell growth was monitored using absorbance at 600 nm taken every 15 minutes using a Tecan Infinite M-plex or Nano.

### Transcription reporter assays

Bioluminescence from each strain was measured in a Tecan Infinite M Plex for 18 hours following 1:200 dilution of stationary phase culture in rich media into freshly prepared minimal media containing the indicated carbon sources in 384-well white microplates with clear bottoms (Corning). Each measurement was adjusted by cell growth using absorbance at 600 nm taken every 15 minutes and normalized to isogenic strains harboring the promoter-less p*Bolux* plasmid as previously described^24^.

### qPCR

Stationary phase cultures were diluted 50-fold into pre-reduced minimal media containing the indicated carbon source and incubated to mid-logarithmic phase (OD_600_ = 0.45 - 0.65). For PMOG grown cultures, glucose or fructose was added to a final concentration of 0.2% and incubated for 10- or 60-minutes, respectively, before 1.0 mL of culture was pelleted by centrifugation, immediately placed on dry ice, and stored at -80°C. RNA was isolated using the RNeasy Mini Kit (QIAGEN), quantified by absorbance, and 1 µg was converted to cDNA following addition of the Superscript IV VILO Master Mix with ezDNAse (Invitrogen) per the manufacturer’s directions. Transcript abundances were measured as previously described^10^ using a QuantStudio5 (ThermoFisher) and PowerUp SYBR Green Master Mix (ThermoFisher) with amplicon specific primers listed in Supplementary Table 1. The 16S *rRNA, rrs,* was used as the reference gene for all qPCR experiments.

### E. coli complementation

Plasmids harboring putative *pfk* genes were introduced into the *pfkAB*-deficient *E. coli* strain as previously described^29^. The resulting strains were cultured in M9 minimal media containing glucose as the sole carbon source supplemented with 100 μM IPTG.

### Protein expression and purification

Proteins were recombinantly expressed as previously described^10^. Briefly, inserts encoding each enzyme were amplified from *B. thetaiotaomicron* VPI-5482 genomic DNA and cloned into pT7-7-N6H4A linearized with NotI-HF and HindIII-HF using NEBuilder Hi-Fi Master Mix (NEB). The resulting plasmids were transformed into chemically competent BL21 cells and plated on selective media. A single colony was inoculated into Luria Bertani broth (BD) and cultured to mid-logarithmic phase before 500 µM IPTG addition and subsequent incubation for 5 hours at 30°C. Pelleted cells were lysed in Lysis Buffer (50 mM sodium phosphate (pH = 7.4) and 300 mM sodium chloride) and N-terminally hexa-histidine-tagged proteins were recovered following co-incubation with Ni^2+^-NTA resin (Thermo) and elution with Lysis Buffer containing 250 mM imidazole. Recovered proteins were buffer exchanged in 10 mM Tris (pH = 7.4) containing 10% glycerol using appropriate molecular weight cut-off centrifugal concentrators (MilliporeSigma). Protein concentrations were determined using a BCA protein quantification kit (Thermo).

### Enzyme assays

Crude lysates and recombinant enzymes were tested by coupling glycolytic reactions with NADH oxidation using an excess of the axillary glycolytic enzymes aldolase (ALD), triose phosphate isomerase (TPI), and glyceraldehyde 3-phosphate dehydrogenase (GDH) (MilliporeSigma). All reactions were performed with 5 µg of enzyme, or 50 µL of lysate, in 100 µL of buffer containing of 50 mM Tris (pH = 7.4), 3 mM MgCl_2_, and 10 mM NH_4_Cl. The degradation of NADH was kinetically measured using absorbance at 340 nm with a Tecan Spark. Changes in absorbance across a five-minute linear slope were converted to moles of NADH consumed using a standard curve and normalized by the amount of enzyme to yield the enzymatic rate (nmol/min/µg protein). Enzyme activities were analyzed using non-linear regression and fit to Michaelis-Menten models using GraphPad Prism. PPi inhibition assays were performed in an identical manner with 5 µg of enzyme, 20 mM substrate, and fit to a dose-response model.

### Quantification of pyrophosphate

*Wild-type Bt* was subbed 1:50 from rich media to minimal media containing the indicated carbon source and grown to mid-logarithmic phase (OD∼0.45 - 0.65). Cells pellets were resuspended in 1 mL buffer containing 50 mM Tris (pH = 7.4), 3 mM MgCl_2_, and 10 mM NH_4_Cl and immediately boiled. PPi was quantified in resulting extracts using a high sensitivity detection kit (MilliporeSigma) according to the manufacturer’s directions.

### Electromobility shift assays

DNA fragments containing the *fusA2* promoter region were amplified by PCR using Q5 High Fidelity Master Mix and isolated from an agarose gel using a QIAquick gel extraction kit (QIAGEN). Equal amounts of probe and 2 mg/mL poly-dI/dC (MilliporeSigma) were combined with purified Cur protein in binding buffer (20 mM HEPES (pH = 8.0), 10 mM KCl, 2 mM MgCl_2_, 0.1 mM EDTA, 0.1 mM dithiothreitol (DTT), and 10% glycerol) in the presence or absence of 1 mM metabolite or 1 µL fractionated *Bt* lysate to a final volume of 20 µL. Reactions were incubated for 20 minutes at room temperature with 4 µL of 6x Native loading buffer, followed by a second, 12-hour incubation at room temperature. Samples containing recombinant Cur, DNA probes, and metabolites or cell extract fractions were then loaded onto a 4-20% TBE gel (Life Technologies), preconditioned in 0.38% TBE, and electrophoresed for 3 hours at 100V. Gels were stained using SYBR green (Invitrogen) per the manufacturer’s directions and imaged using an AI680 (GE).

### Separation of soluble metabolites by liquid chromatography

*Wild-type Bt* cell pellets were lysed in 100-fold lower volume of phosphate buffered saline (MilliporeSigma) by bead beating for five 40-second intervals in a pre-chilled aluminum block interspersed with 5-minute incubations at 4°C, followed by centrifugation for 5 minutes at 18,213x*g*. The supernatant was recovered and fractionated using an Agilent Infinity 1260 fitted with a ZORBAX StableBond C18 Column (Agilent). The injection volume was set to 100 µL and samples were eluted using a linear 20 minute gradient (water:MeOH) with a flow rate of 0.5 mL/min. Fractions were dried under air and resuspended in 50 µL of ultrapure water for use in EMSA assays.

### Metabolomics

*Bt* strains were cultured in minimal media containing 5 mg/mL glucose to mid-logarithmic phase (OD∼0.45 - 0.65) where 0.5 ODs were collected by centrifugation for 30 seconds, quickly decanted, and immediately flash frozen in a mixture of ethanol and dry ice. Metabolite abundances were measured using Targeted Quantification Analysis by LC-MRM/MS at Creative Proteomics.

### In vivo competitive fitness of B. thetaiotaomicron strains

All animal experiments were performed in accordance with protocols approved by Penn State Institutional Animal Care and Use Committee. Germ-free C57/BL6 mice were maintained in flexible plastic gnotobiotic isolators with a 12-hour light/dark cycle and provided a standard, autoclaved mouse chow (LabDiet, 5021) or high-sugar chow (Bioserv, S4944) *ad libitum^10^*. Mice were gavaged with 10^8^ CFU of each indicated strains suspended in 200 μL of phosphate-buffered saline. Input (day 0) abundance of each strain was determined by CFU plating. Fecal pellets were collected at the desired times and genomic DNA was extracted as described previously^12^. The abundance of each strain was measured by qPCR, using barcode-specific primers (Supplemental Table 2) as described previously^10^.

### Software and statistics

Data collection and curation was done in Microsoft Excel. All data was plotted in GraphPad Prism, except for the heatmap in Fig. 4, which was generated using ggplot2 in RStudio. Statistical analyses were also calculated in Prism. For all experiments, *n* denotes individual biological replicates across two independent experiments.

## Acknowledgements

We thank Jordan Bisanz, Donalee McElrath, Michelle Irish, Jacob Perryman, and Alexandra Sprinkle for assistance with germ-free mouse experiments. Additionally, we thank Andrew Goodman, Whitman Schofield, Andrew Patterson, and Imhoi Koo for insights about metabolite measurements. Finally, we thank Jennifer Lausch and Jessica Gaydos for helpful comments during the preparation of this manuscript. This work was supported, in part, by National Institutes of Health Grants DK132711 and GM147178 to G.E.T. and GM123798 to E.A.G.

## Notes

### Competing Interest Statement

The authors have declared no competing interest.

### Summary of Updates

New data were added demonstrating that the hexoses fucose and galNAc do not elicit Cur inhibition like glucose and fructose. Furthermore, the text has been revised to more clearly indicate genes within the Bt fructan PUL. Finally, information was added into the methods section to provide a comprehensive description of all experimental approaches described in the manuscript.

