## Extended Data Fig. 1 for "Hierarchical glycolytic pathways control the carbohydrate utilization regulator in human gut *Bacteroides*"

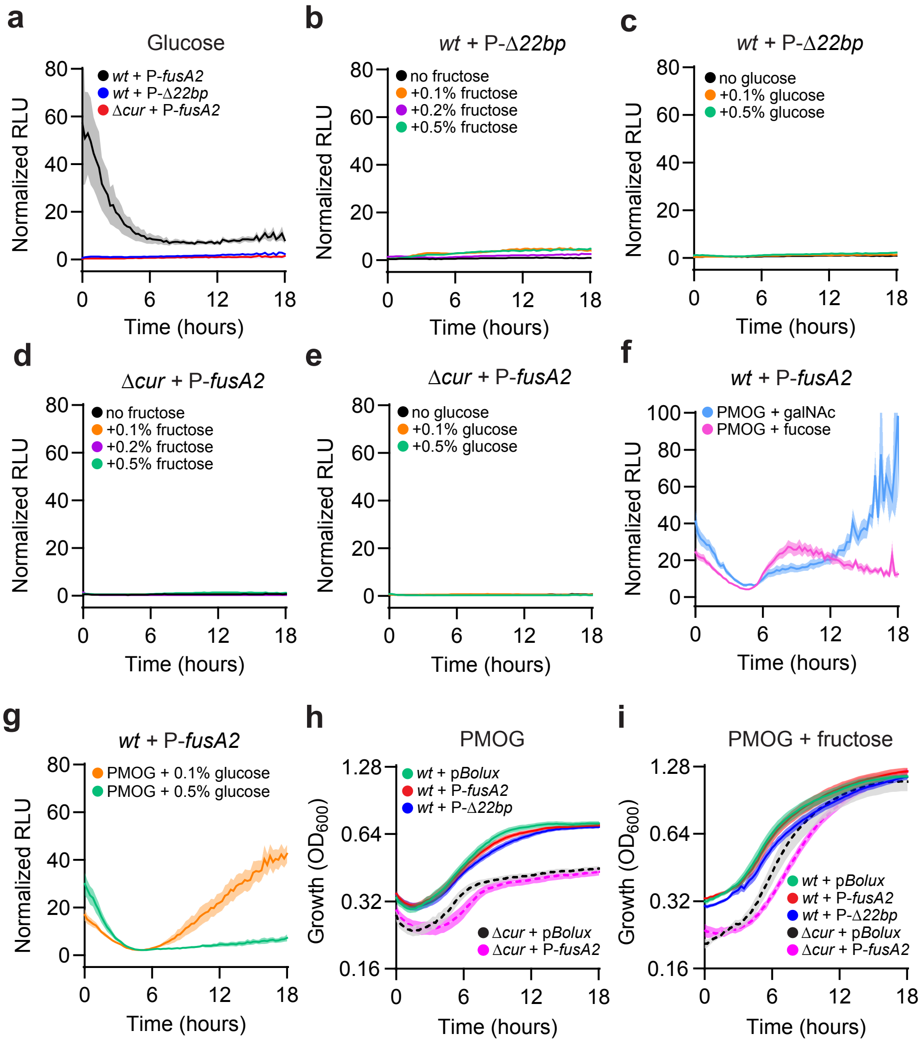


**Extended Data Fig. 1. P-*fusA2* bioluminescence requires *cur*.** **a,** Normalized bioluminescence from a *wild-type* strain harboring P-*fusA2* (black) or P-Δ*22bp* (blue) and a Δ*cur* strain harboring P-*fusA2* (red) cultured in glucose as a sole carbon source. **b,** Normalized bioluminescence from *wild-type* harboring P-Δ*22bp* cultured in PMOG alone (black) or in combination with 0.1% (orange), 0.2% (purple), or 0.5% (green) fructose. **c,** Normalized bioluminescence from *wild-type* harboring P-Δ*22bp* cultured in PMOG alone (black) or in combination with 0.1% (orange) or 0.5% (green) glucose. **d,** Normalized bioluminescence from Δ*cur* harboring P-*fusA2* cultured in PMOG alone (black) or in combination with 0.1% (orange), 0.2% (purple), or 0.5% (green) fructose. **e,** Normalized bioluminescence from Δ*cur* harboring P-*fusA2* cultured in PMOG alone (black) or in combination with 0.1% (orange) or 0.5% (green) glucose. **f,** Normalized bioluminesence of *wild-type* harboring P-*fusA2* cultured in PMOG mixtures containing galNAc (light blue) or fucose (pink). **g,** Normalized bioluminescence from *wild-type* harboring P-*fusA2* cultured in PMOG with 0.1% (orange) or 0.5% (green) glucose. **h,i,** Growth of *wild-type* (solid) harboring p*Bolux* (green), P-*fusA2* (red), or P-Δ*22bp* (blue) and Δ*cur* (dashed) harboring p*Bolux* (black) or P-*fusA2* (fushia) grown in **(h)** PMOG alone or **(i)** equal amounts of PMOG and fructose. For all panels, n=8; error is SEM in color matched shading.
