## Extended Data Fig. 2 for "Hierarchical glycolytic pathways control the carbohydrate utilization regulator in human gut *Bacteroides*"

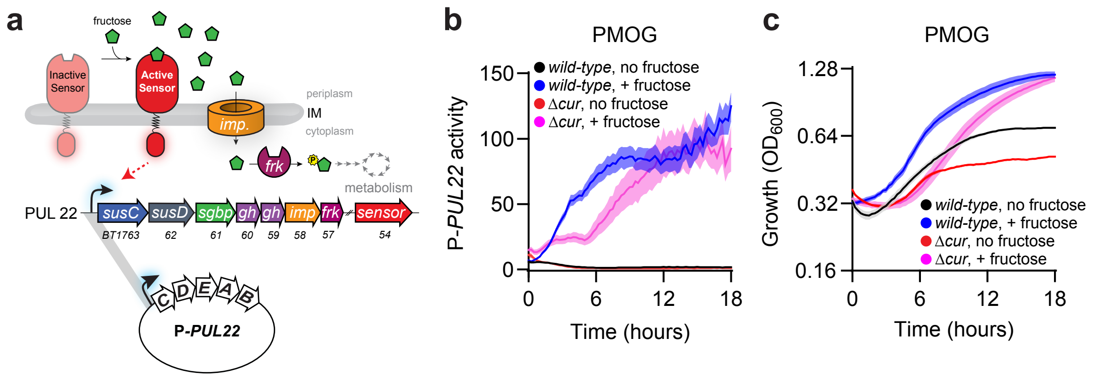


**Extended Data Fig. 2. Cur is dispensable for fructose-inducible bioluminescence. a,** Schematic depicting the fructan utilization PUL (PUL22). Sensor$^{PUL22}$ binds fructose in the periplasm and subsequently activates transcription of *PUL22* in the cytoplasm. The plasmid P-*PUL22* harbors the sensor$^{PUL22}$-dependent promoter preceding *susC$^{PUL22}$* and confers fructose-responsive bioluminescence. **b,** Normalized bioluminescence from *wild-type* or Δ*cur* harboring P-*PUL22* were cultured in PMOG as the sole carbon source (black & red, respectively) or in combination with 0.5% fructose (blue & pink, respectively). **c,** Growth of strains described in **(b)** in an equal mixture of 0.5% PMOG and fructose. For panel **a,** abbreviations are as follows: inner membrane (IM), surface glycan binding protein (*sgbp*), glycosyl hydrolase (*gh*), sugar importer (*imp*), fructokinase (*frk*). For panels **b,c,** n=8; error is SEM in color matched shading.
