## Extended Data Fig. 3 for "Hierarchical glycolytic pathways control the carbohydrate utilization regulator in human gut *Bacteroides*"

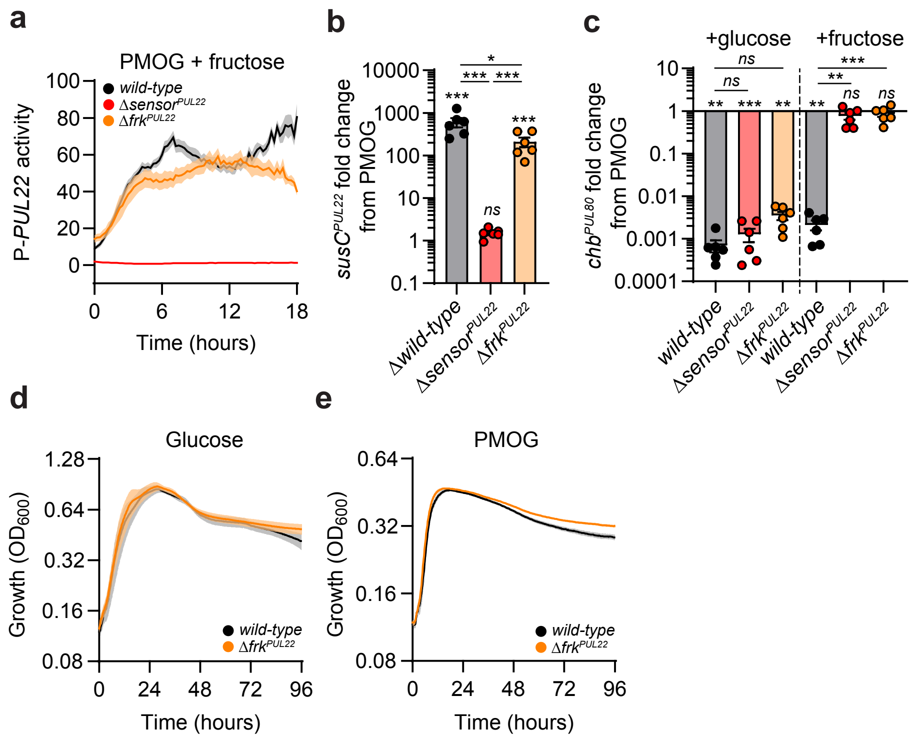


**Extended Data Fig. 3. Fructose phosphorylation is dispensable for *sensor*$^{PUL22}$-dependent PUL activation but required for Cur inhibition. a,** Normalized bioluminescence from *wild-type* (black), Δ*sensor*$^{PUL22}$ (red), or Δ*frk*$^{PUL22}$ (orange) harboring P-*PUL22* cultured in media containing equal amounts of PMOG and fructose. **b,** Fold change of *susC*$^{PUL22}$ transcript amounts from *wild-type*, Δ*sensor*$^{PUL22}$, or Δ*frk*$^{PUL22}$ at 60 minutes following the addition of 0.2% fructose to cells cultured in media containing PMOG as the sole carbon source. **c,** Fold change of *chb*$^{PUL80}$ transcript amounts from *wild-type*, Δ*sensor*$^{PUL22}$, or Δ*frk*$^{PUL22}$ following the addition of 0.2% glucose (left) or fructose (right). **d,e,** Growth of *wild-type* or Δ*frk*$^{PUL22}$ in **(d)** glucose or **(e)** PMOG as the sole carbon source. For panels **a,d,e,** n=8; error is SEM in color matched shading. For panel **b,** n=6; error is SEM; *P*-values were calculated by 1-way ANOVA with Fisher's LSD test. For panel **c,** n=6; error is SEM; *P*-values were calculated by 2-way ANOVA with Fisher's LSD test. For panels **b,c,** * represents values < 0.05, ** < 0.01, *** < 0.001.
