## Extended Data Fig. 4 for "Hierarchical glycolytic pathways control the carbohydrate utilization regulator in human gut *Bacteroides*"

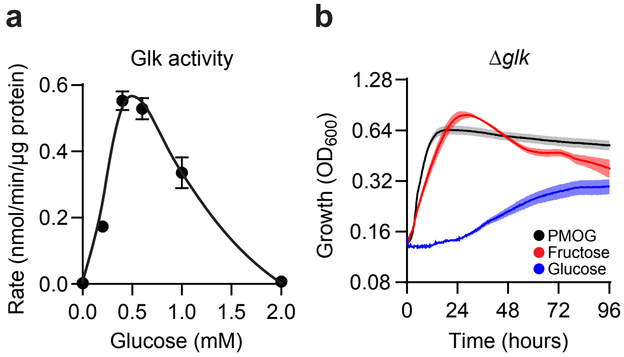


**Extended Data Fig. 4. Glk facilitates glucose phosphorylation and utilization. a,** Glucokinase activity of purified Glk protein. **b,** Growth of Δ*glk* in PMOG (black), fructose (red), or glucose (blue) as sole carbon sources. For panel **a**, n=4; error is SEM. For panel **b**, n=8; error is SEM with color matched shading.
