## Extended Data Fig. 5 for "Hierarchical glycolytic pathways control the carbohydrate utilization regulator in human gut *Bacteroides*"

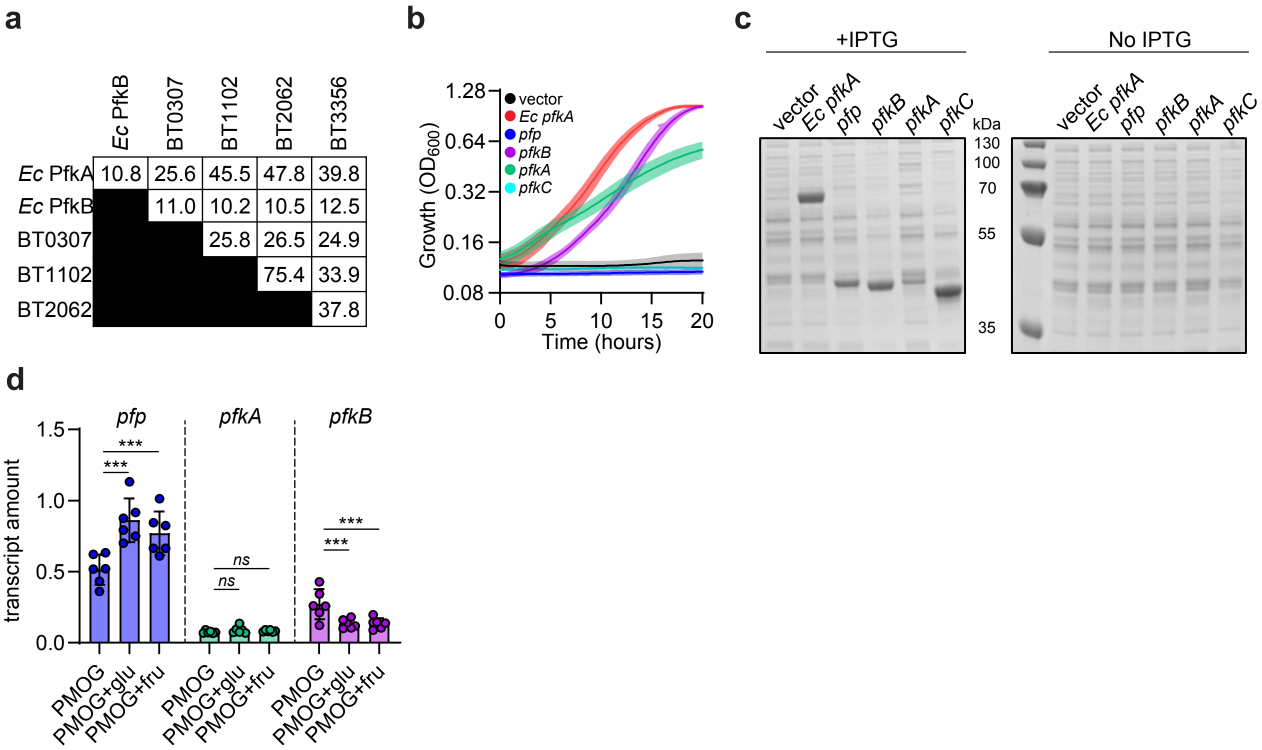


**Extended Data Fig. 5. *Bt* possesses 2 *bona fide* Pfk enzymes. a,** Amino acid sequence identity between *Ec* PfkA and PfkB with putative fructose bisphosphate biosynthetic enzymes in *Bt*. **b,** Growth of Δ*pfkAB Ec* harboring an empty vector (black), or a plasmid encoding *Ec pfkA* (red), *pfp* (blue), *pfkB* (purple), *pfkA* (green) or *pfkC* (cyan) in minimal media containing glucose and 100 µM IPTG. **c,** Total protein from strains described in **(b)** grown in rich media with (left) or without (right) the addition of 100 µM IPTG. **d,** qPCR analysis of *pfp*, *pfkA*, and *pfkB* transcript amounts from *wild-type* grown in minimal media containing PMOG and 60 minutes following the addition of 0.2% glucose or fructose, respectively. For panel **b,** n=8; error is SEM with color matched shading. For panel **d,** n=6; error is SEM; *P*-values were calculated by 1-way ANOVA with Fisher's LSD test and * represents values < 0.05, ** < 0.01, *** < 0.001.
