## Extended Data Fig. 6 for "Hierarchical glycolytic pathways control the carbohydrate utilization regulator in human gut *Bacteroides*"

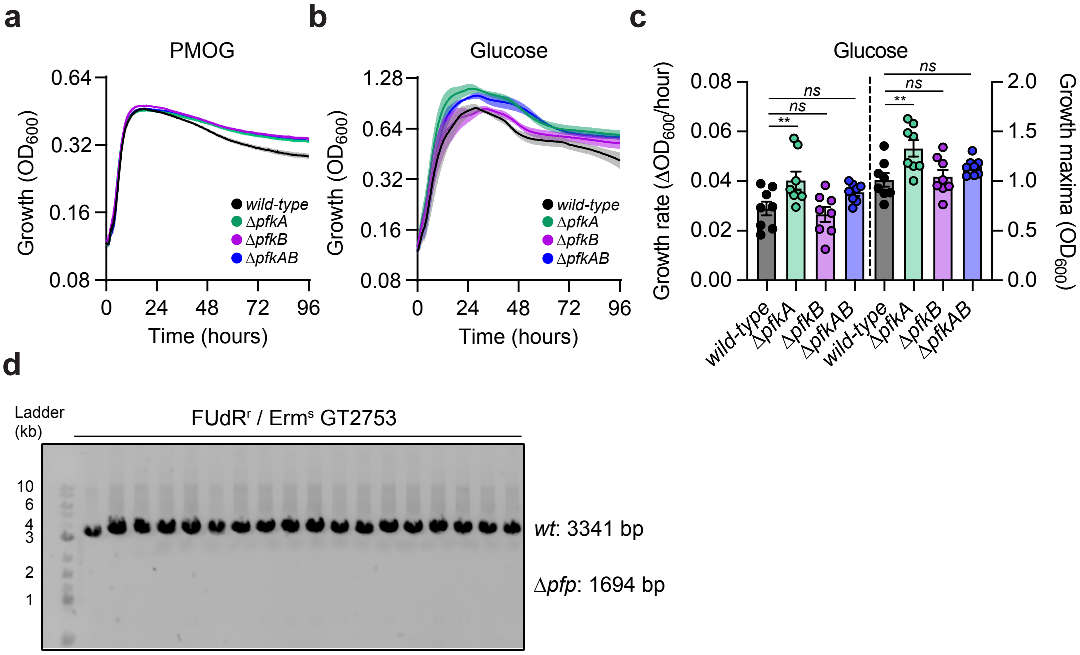


**Extended Data Fig. 6. Phosphofructokinases are dispensable for growth. a,b,** Growth of *wild-type* (black), Δ*pfkA* (green), Δ*pfkB* (purple), or Δ*pfkAB* (blue) in minimal media containing **(a)** PMOG or **(b)** glucose as the sole carbon source. **c,** Growth rate and maxima of *wild-type*, Δ*pfkA*, Δ*pfkB*, or Δ*pfkAB* grown in glucose as the sole carbon source. **d,** Putative *pfp* deletions examined by PCR analysis. For panels **a,b,** n=8; error is SEM with color matched shading. For panel **c,** n=8, error is SEM. P-values were calculated using 1-way ANOVA with Fisher's LSD test and ** represents values < 0.01.
