## Extended Data Fig. 7 for "Hierarchical glycolytic pathways control the carbohydrate utilization regulator in human gut *Bacteroides*"

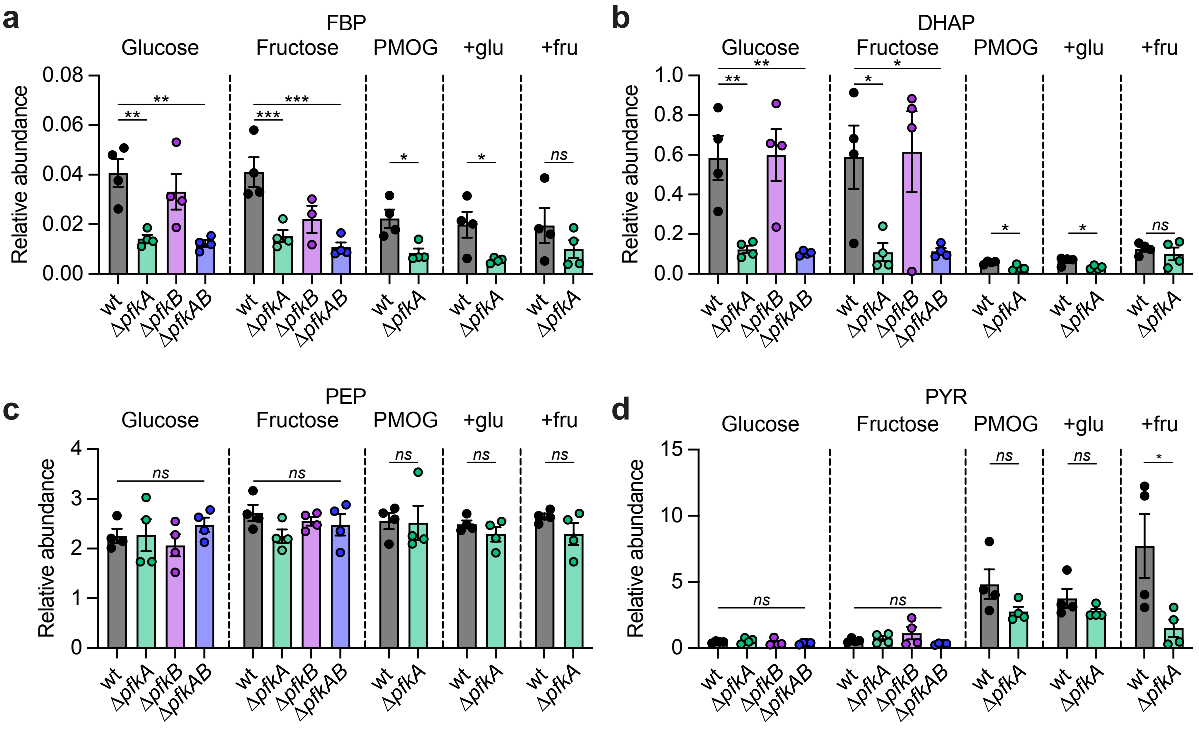


**Extended Data Fig. 7. Relative abundance of glycolytic metabolites in *pfk* mutants. a-d,** Relative amounts of **(a)** fructose bisphosphate (FBP), **(b)** dihydroxyacetone-P (DHAP), **(c)** phosphoenolpyruvate (PEP), or **(d)** pyruvate (PYR) from *wild-type*, Δ *pfkA*, Δ*pfkB*, or Δ*pfkAB* grown in minimal media containing glucose, fructose, PMOG, PMOG with glucose (+glu), or PMOG with fructose (+fru) as a sole carbon source. For all panels, n=4; error is SEM. For glucose and fructose**,** *P*-values were calculated using 2-way ANOVA with Fisher's LSD test and * represents values < 0.05, ** < 0.01, *** < 0.001. For PMOG, PMOG with glucose, or PMOG with fructose, *P*-values were calculated using an unpaired t-test and * represents values < 0.05.
