## Extended Data Fig. 8 for "Hierarchical glycolytic pathways control the carbohydrate utilization regulator in human gut *Bacteroides*"

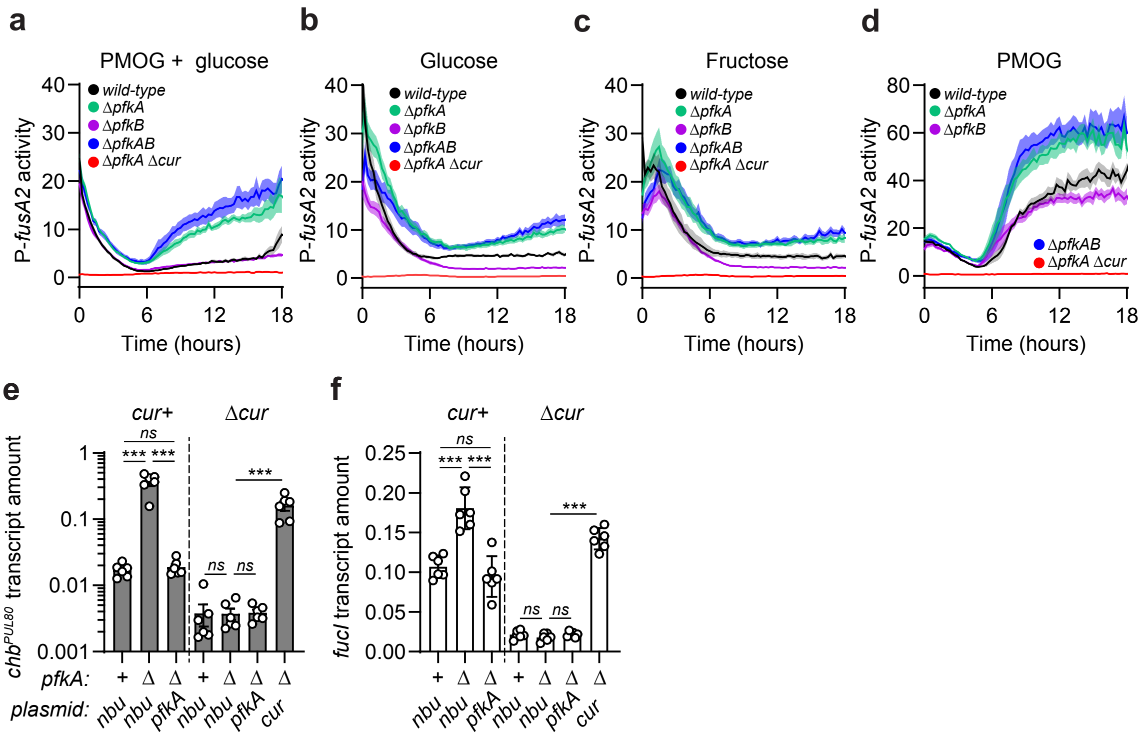


**Extended Data Fig. 8. ATP-dependent FBP production silences Cur *in vitro*. a-d,** Normalized bioluminescence from *wild-type* (black)*,* Δ*pfkA* (green), Δ*pfkB* (purple), Δ*pfkAB* (blue), or Δ*pfkA* Δ*cur* (red) harboring P-*fusA2* grown in minimal media containing equal amounts of **(a)** PMOG and glucose, **(b)** glucose, **(c)** fructose, or **(d)** PMOG as a sole carbon source. **e,f,** Transcript amounts of **(e)** *chb*$^{PUL80}$ or **(f)** *fucI* in *wild-type*, Δ*pfkA*, Δ*cur*, or Δ*pfkA* Δ*cur* harboring empty vector (*nbu*) or complementing plasmids grown in glucose as the sole carbon source. For panels **a-d,** n=8; error is SEM with color matched shading. For panels **e,f,** *P*-values were calculated by 2-way ANOVA with Fisher's LSD test and *** represents values < 0.001.
