## Extended Data Fig. 9 for "Hierarchical glycolytic pathways control the carbohydrate utilization regulator in human gut *Bacteroides*"

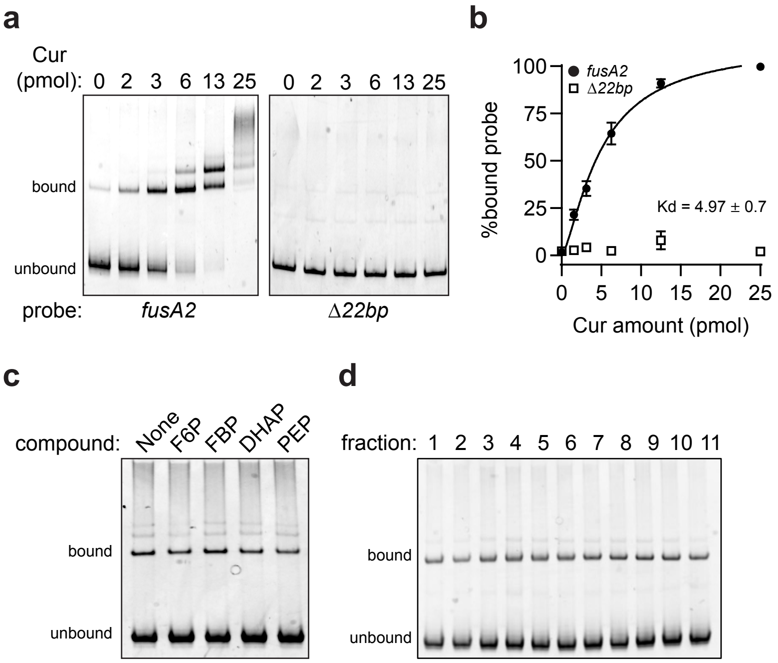


**Extended Data Fig. 9. FBP does not alter *in vitro* Cur binding to the *fusA2* promoter. a,** EMSA containing a *wild-type* or Δ*22bp fusA2* promoter DNA fragment co-incubated with the indicated amounts of Cur protein. **b,** DNA-binding saturation curve of **(a)**. **c,** EMSA assay containing 3 pmol of Cur protein incubated with the *fusA2* promoter fragment and 1 mM of glycolytic intermediates. **d,** EMSA assay containing 3 pmol of Cur protein with fractionated *wild-type Bt* lysate. For panel **b,** n=3; error is SEM.
