## Extended Data Fig. 10 for "Hierarchical glycolytic pathways control the carbohydrate utilization regulator in human gut *Bacteroides*"

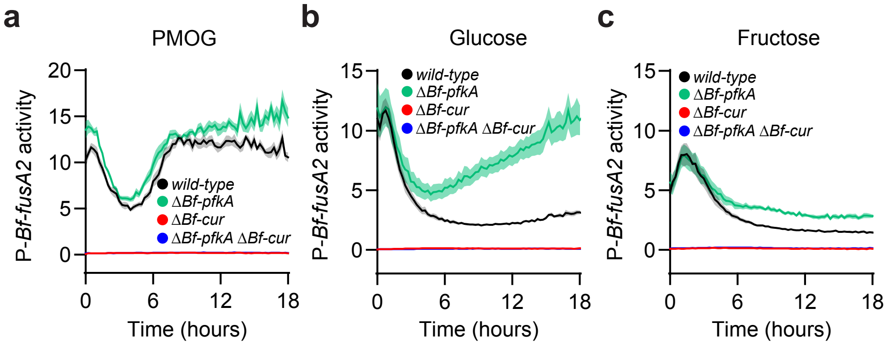


**Extended Data Fig. 10. *Bf-pfkA* is required for Cur inhibition in *B. fragilis*. a-c,** Normalized bioluminescence from *wild-type* (black), Δ*Bf-pfkA* (green), Δ*Bf-cur* (red), or Δ*Bf-pfkA* Δ*Bf-cur* (blue) harboring P-*Bf-fusA2* cultured in media containing **(a)** PMOG, **(b)** glucose, or **(c)** fructose as the sole carbon source. For all panels, n=8; error is SEM in color matched shading.
