## Extended Data Tables 1&2 for "Hierarchical glycolytic pathways control the carbohydrate utilization regulator in human gut *Bacteroides*"

Supplementary Table 1. Strains and plasmids used in this study

| **Identifier** | ***Description*** | **Genotype** | **Source** |
| --- | --- | --- | --- |
| ***B. thetaiotaomicron*** | | | |
| GT23 | *A tdk-deficient strain considered wild-type in this work* | *Δtdk* | ^1^ |
| WH311 | *A strain lacking the 22bp Cur binding site preceding fusA2* | *Δtdk Δ22bp* | ^2^ |
| GT593 | *A strain harboring an in-frame C-terminally HA-tagged roc* | *∆tdk BT3172-HA* | ^3^ |
| GT1867 | *A wild-type strain harboring pBolux* | *Δtdk + pBolux* | ^4^ |
| GT4316 | *A wild-type strain harboring P-fusA2* | *Δtdk + P-fusA2* | This study |
| GT4317 | *A wild-type strain harboring P-Δ22bp* | *Δtdk + P-Δ22bp* | This study |
| GT1979 | *A cur-deficient strain harboring pBolux* | *Δtdk ΔBT4338 + pBolux* | This study |
| GT4318 | *A cur-deficient strain harboring P-fusA2* | *Δtdk ΔBT4338 + P-fusA2* | This study |
| GT1893 | *A wild-type strain harboring P-PUL22* | *Δtdk +* P-*BT1763* | ^4^ |
| GT2000 | *A cur-deficient strain harboring P-PUL22* | *Δtdk ΔBT4338 +* P-*BT1763* | This study |
| GT165 | *A sensor^PUL22^-deficient strain* | *Δtdk ΔBT1754* | ^4^ |
| GT2619 | *A sensor^PUL22^-deficient strain harboring pBolux* | *Δtdk ΔBT1754 + pBolux* | ^4^ |
| GT2620 | *A sensor^PUL22^-deficient strain harboring P-PUL22* | *Δtdk ΔBT1754 + P-BT1763* | ^4^ |
| GT953 | *A frk^PUL22^-deficient strain* | *Δtdk ΔBT1757* | This study |
| GT4527 | *A frk^PUL22^-deficient strain harboring pBolux* | *Δtdk ΔBT1757 + pBolux* | This study |
| GT4528 | *A frk^PUL22^-deficient strain harboring P-PUL22* | *Δtdk ΔBT1757 + P-BT1763* | This study |
| GT4529 | *A frk^PUL22^-deficient strain harboring P-fusA2* | *Δtdk ΔBT1757 + P-fusA2* | This study |
| GT4817 | *A glk-deficient strain* | *Δtdk ΔBT2493* | This study |
| GT4878 | *A glk-deficient strain harboring pBolux* | *Δtdk ΔBT2493* + *pBolux* | This study |
| GT4879 | *A glk-deficient strain harboring P-fusA2* | *Δtdk ΔBT2493* + *P-fusA2* | This study |
| GT758 | *A pfkA-deficient strain* | *Δtdk ΔBT2062* | This study |
| GT945 | *A pfkB-deficient strain* | *Δtdk ΔBT1102* | This study |
| GT949 | *A pfkA and pfkB-deficient strain* | *Δtdk ΔBT2062*  *Δtdk ΔBT1102* | This study |
| GT2753 | *A strain harboring pEXCHANGE-ΔBT0307 integrated immediately upstream of BT0307* |  |  |
| GT4464 | *A pfkA-deficient strain harboring pBolux* | *Δtdk ΔBT2062 + pBolux* | This study |
| GT4471 | *A pfkA-deficient strain harboring P-fusA2* | *Δtdk ΔBT2062 + P-fusA2* | This study |
| GT4466 | *A pfkB-deficient strain harboring pBolux* | *Δtdk ΔBT1102 + pBolux* | This study |
| GT4473 | *A pfkB-deficient strain harboring P-fusA2* | *Δtdk ΔBT1102 + P-fusA2* | This study |
| GT4612 | *A pfkA and pfkB-deficient strain harboring pBolux* | *Δtdk ΔBT2062 ΔBT1102 + pBolux* | This study |
| GT4613 | *A pfkA and pfkB-deficient strain harboring P-fusA2* | *Δtdk ΔBT2062 ΔBT1102 + P-fusA2* | This study |
| GT1313 | *A cur and pfkA-deficient strain* | *Δtdk ΔBT4338 ΔBT2062* | This study |
| GT4887 | *A cur and pfkA-deficient strain harboring pBolux* | *Δtdk ΔBT4338 ΔBT2062 + pBolux* | This study |
| GT4888 | *A cur and pfkA-deficient strain harboring P-fusA2* | *Δtdk ΔBT4338 ΔBT2062 + P-fusA2* | This study |
| GT3164 | *A wild-type strain harboring pNBU2-tetQb* | *Δtdk + pNBU2-tetQb* | This study |
| GT4704 | *A pfkA-deficient strain harboring pNBU2-tetQb* | *Δtdk ΔBT2062*  *+ pNBU2-tetQb* | This study |
| GT4705 | *A pfkA-deficient strain harboring pNBU2-pfkA* | *Δtdk ΔBT2062*  *+ pNBU2-BT2062* | This study |
| GT1339 | *A cur-deficient strain harboring pNBU2-tetQb* | *Δtdk ΔBT4338*  *+ pNBU2-tetQb* | This study |
| GT4706 | *A cur- and pfkA-deficient strain harboring pNBU2-tetQb* | *Δtdk ΔBT4338 ΔBT2062*  *+ pNBU2-tetQb* | This study |
| GT4707 | *A cur- and pfkA-deficient strain harboring pNBU2-pfkA* | *Δtdk ΔBT4338 ΔBT2062*  *+ pNBU2-BT2062* | This study |
| GT4708 | *A cur- and pfkA-deficient strain harboring pNBU2-cur* | *Δtdk ΔBT4338 ΔBT2062*  *+ pNBU2-BT4338* | This study |
| GT4702 | *A wild-type strain harboring pNBU2-BC-01* | *Δtdk + pNBU2-BC-01* | This study |
| GT4698 | *A pfkA-deficient strain harboring pNBU2-BC-04* | *Δtdk ΔBT2062 + pNBU2-BC-04* | This study |
| GT4703 | *A cur-deficient strain harboring pNBU2-BC-01* | *Δtdk ΔBT4338 + pNBU2-BC-01* | This study |
| GT4700 | *A cur- and pfkA-deficient strain harboring pNBU2-BC-04* | *Δtdk ΔBT4338 ΔBT2062 + pNBU2-BC-04* | This study |
| ***B. fragilis*** | | | |
| *ATCC 25285* | *Wild-type Bf* |  | ATCC |
| GT2520 | *A Bf-cur-deficient strain* | *ΔBF9343_0915* | ^5^ |
| GT4813 | *A Bf-pfkA-deficient strain* | *ΔBF9343_3444* | This study |
| GT4855 | *A Bf-cur and Bf-pfkA-deficient strain* | *ΔBF9343_0915 ΔBF9343_3444* | This study |
| GT3428 | *Wild-type Bf harboring pBolux* | *Wild-type + pBolux* | This study |
| GT4571 | *Wild-type Bf harboring P-Bf-fusA2* | *Wild-type + P-Bf-fusA2* | This study |
| GT4669 | *A Bf-cur-deficient strain harboring pBolux* | *ΔBF9343_0915 + pBolux* | This study |
| GT4693 | *A Bf-cur-deficient strain harboring P-Bf-fusA2* | *ΔBF9343_0915 + P-Bf-fusA2* | This study |
| GT4864 | *A Bf- pfkA-deficient strain harboring pBolux* | *ΔBF9343_3444 + pBolux* | This study |
| GT4867 | *A Bf-pfkA-deficient strain harboring P-Bf-fusA2* | *ΔBF9343_3444 + P-Bf-fusA2* | This study |
| GT4868 | *A Bf-cur and Bf-pfkA-deficient strain harboring pBolux* | *ΔBF9343_0915 ΔBF9343_3444 + pBolux* | This study |
| GT4871 | *A Bf-cur and Bf-pfkA-deficient strain harboring P-Bf-fusA2* | *ΔBF9343_0915 ΔBF9343_3444 + P-Bf-fusA2* | This study |
| ***E. coli*** |  |  |  |
| GT1 | *S17-1* | *λpir* | ^6^ |
| BL21 (DE3) | *A strain for engineering protein over-expression* | *lon-11 Δ(ompT-nfrA)885 Δ(galM-ybhJ)884 λDE3 Δ46 [mal+]K-12(λ^S^) hsdS10* | ^7^ |
| RL257 | *A pfk-deficient strain* | *F-, [araD139]B/r, lacIp-4000(lacIQ), e14-, pfkB205(del-ins)-FRT, flhD5301, Δ(fruK-yeiR)725(fruA25), relA1, rpsL150(strR), rbsR22, pfkA203(del-ins)-FRT, Δ(fimB-fimE)632(-IS1), deoC1* | ^8^ |
| GT3382 | *A pfk-deficient strain harboring an empty vector* | *RL257 + pUHE21-lacIQ* | This study |
| GT3383 | *A pfk-deficient strain harboring an IPTG-inducible BT0307* | *RL257 + pUHE21-lacIQ-BT0307* | This study |
| GT3384 | *A pfk-deficient strain harboring an IPTG-inducible BT1102* | *RL257 + pUHE21-lacIQ-BT1102* | This study |
| GT3385 | *A pfk-deficient strain harboring an IPTG-inducible BT2062* | *RL257 + pUHE21-lacIQ-BT2062* | This study |
| GT3386 | *A pfk-deficient strain harboring an IPTG-inducible BT3356* | *RL257 + pUHE21-lacIQ-BT3356* | This study |
| GT4231 | *A pfk-deficient strain harboring an IPTG-inducible Ec pfkA* | *RL257 + pUHE21-lacIQ-pfkA* | This study |
| **Plasmids** | | | |
| *Name* | *Description* | | *Source* |
| p*Bolux* | A *Bacteroides*-optimized bioluminescent reporter plasmid | | **^4^** |
| P-*fusA2* | A reporter of *BT2167* expression | | This study |
| pUHE21-lacIQ | A vector for IPTG-inducible gene expression | |  |
| pUHE21-*lacIQ*-*pfkA* (*Ec*) | Facilitate IPTG-inducible BT1102 expression | | This study |
| pUHE21-*lacIQ-BT1102* | Facilitate IPTG-inducible BT1102 expression | | This study |
| pUHE21-*lacIQ-BT2062* | Facilitate IPTG-inducible BT2062 expression | | This study |
| pUHE21-*lacIQ-BT3356* | Facilitate IPTG-inducible BT3356 expression | | This study |
| pUHE21-*lacIQ-BT0307* | Facilitate IPTG-inducible BT0307 expression | | This study |
| pT7-7-N-6H-4A | A plasmid for engineering T7-promoted products with an N-terminal hexa-histidine tag | | This study |
| pT7-7-N-6H-4A-BT1757 | Express N-terminally hexa-histidine tagged BT1102 | | This study |
| pT7-7-N-6H-4A-BT2493 | Express N-terminally hexa-histidine tagged BT1102 | | This study |
| pT7-7-N-6H-4A-BT1102 | Express N-terminally hexa-histidine tagged BT1102 | | This study |
| pT7-7-N-6H-4A-BT2062 | Express N-terminally hexa-histidine tagged BT2062 | | This study |
| pT7-7-N-6H-4A-BT3356 | Express N-terminally hexa-histidine tagged BT3356 | | This study |
| pT7-7-N-6H-4A-BT0307 | Express N-terminally hexa-histidine tagged BT0307 | | This study |
| pT7-7-BT4338-4xG-6xH | Express C-terminally hexa-histidine tagged BT4338 | | This study |
| pEXCHANGE*-tdk* | A plasmid to engineer deletions by allelic exchange in *Bt* | | ^1^ |
| pEXCHANGE-*ΔBT1757* | A plasmid to engineer a deletion of *BT1757* | | This study |
| pEXCHANGE-*ΔBT2493* | A plasmid to engineer a deletion of *BT2493* | | This study |
| pEXCHANGE-*ΔBT4338* | A plasmid to engineer a deletion of *BT4338* | | ^9^ |
| pEXCHANGE-*ΔBT2062* | A plasmid to engineer a deletion of *BT2062* | | This study |
| pEXCHANGE-*ΔBT1102* | A plasmid to engineer a deletion of *BT1102* | | This study |
| pEXCHANGE-*ΔBT0307* | A plasmid to engineer a deletion of *BT0307* | | This study |
| P-*Bf-fusA2* | A reporter of *BF9343_3536* expression | | This study |
| pLGB13 | A plasmid to engineer deletions by allelic exchange in *Bf*. | | ^10^ |
| pLGB13-*ΔBF9343_3444* | A plasmid to engineer deletion of *Bf* *pfkA* | | This study |
| pLGB13-*ΔBF9343_0915* | A plasmid to engineer deletion of *Bf* *cur* | | ^5^ |

Supplementary Table 2. Primers used in this study

| **Identifier** | **Description** | **Sequence** |
| --- | --- | --- |
| **Reporter plasmids** | | |
| pBolux-P-*fusA2*f | Clone the *Bt fusA2* promoter into p*Bolux* | AGCTCGGTACCCGGGGATCCAACGTGTCTTACAGAAGGGAATATG |
| pBolux-P-*fusA2*r |  | AATGCGGGAGTGACTAGTAATATATAAGATTTTAGTAATTACTCAGTATGTTTCTCGC |
| pBolux-P-*Bf*-*fusA2*f | Clone the *Bf fusA2* promoter into p*Bolux* | AGCTCGGTACCCGGGGATCCGAAAAACCTCCGTTTTGCACCG |
| pBolux-P-*Bf-fusA2*r |  | ATGCGGGAGTGACTAGTGATATATAAGATTTAAATGAATGATTCAGTATGTG |
| **qPCR primers** | | |
| qBt16sF | Measuring *16s* rRNA transcript levels from *Bt* by qPCR | GGTAGTCCACACAGTAAACGATGAA |
| qBt16sR |  | CCCGTCAAATTCCTTTGAGTTTC |
| q*BT2167*f | Measure the *Bt fusA2* transcript | AAAACGTCGCGGATCTGTTG |
| q*BT2167*r |  | TGGAGAACGGTAGAGAAAACGG |
| q*BT4295*f | Measure the *BT4295* transcript | AGACCAATGTACCACTCGACAG |
| q*BT4295*r |  | ACTTTGCCCTCGTGTTGTAC |
| q*BT1273*f | Measure the *fucI* transcript | AGACCAATGTACCACTCGACAG |
| q*BT1273*r |  | ACTTTGCCCTCGTGTTGTAC |
| pNBU2_tet_BC01 | Measure relative abundance of *in vivo* competition strains | ATGTCGCCAATTGTCACTTTCTCA |
| pNBU2_tet_BC04 |  | CTCCATAAAGGCGCATACCGACTA |
| UNIV-R |  | CACAATATGAGCAACAAGGAATCC |
| **Deletion primers** | | |
| pEXCHANGE-dBT1757_5f | Engineer a pEXCHANGE plasmid to delete *BT1757* | GCTCTAGAACTAGTGGATCCTTGCCATGTGGGGAGCAACG |
| dBT1757_5r |  | GATCTACAGATTTAATTGTTTTCCAAATATAAA |
| dBT1757_3f |  | AACAATTAAATCTGTAGATCAATCTCTTTAAATTCAAAAAGAAGGTGAAT |
| pEXCHANGE-dBT1757_3r |  | AATAACTTTATGCTTGATACTTTTTGCTTGGTCGACTCGAATGTTATCTT |
| pEXCHANGE-dBT2493_5f | Construct a plasmid to delete *BT2493* using allelic exchange. | GCTCTAGAACTAGTGGATCCATTCCCATTCGATGCAAGTCTTTC |
| dBT2493_5r |  | GATACTTTATTTAAAAGGGTTAGTTAATTATAAAGTTTACTGCACA |
| dBT2493_3f |  | CTAACCCTTTTAAATAAAGTATCTATCATTTTTTTAATAATAAAGGTAATTGGATCCCCG |
| pEXCHANGE-dBT2493_3r |  | AGATAACATTCGAGTCGACAACAACGACCTCCGCATCCA |
| pEXCHANGE-dBT2062_5f | Construct a plasmid to delete *BT2062* using allelic exchange. | GCGGCCGCTCTAGAACTAGTGGATCCGATCGCCCACAAGAATAATATCG |
| dBT2062_5r |  | ATATGAATGTATTACCTTTAAACAATCTTTCATTAATTAGTTTGACGTG |
| dBT2062_3f |  | TAAAGGTAATACATTCATATAAGGTAATATAAAGAACG |
| pEXCHANGE-dBT2062_3r |  | GAAAGAAGATAACATTCGAGTCGACTTTGCAGGATGGAAAGCCGAA |
| pEXCHANGE-dBT1102_5f | Construct a plasmid to delete *BT1102* using allelic exchange. | GCTCTAGAACTAGTGGATCCTTCCAGCAGAGTATCCTGCTGC |
| dBT1102_5r |  | CTGACATATTAACGGTTATTAATTTGTTTTTTATATTTTAATTAATCAGGC |
| dBT1102_3f |  | AATAACCGTTAATATGTCAGAAGTTGCTCC |
| pEXCHANGE-dBT1102_3r |  | AAGATAACATTCGAGTCGACAATGCGGCAGAAAATCCCGC |
| pLGB13-dBF9343_3444-5f | Construct a plasmid to delete *BF9343_3444* using allelic exchange. | TTAGCATTATGAGTGGATCCCTTAACCACACGAAGAGCTTCTTTATC |
| pLGB13-dBF9343_3444-5r |  | AACAATCATGTTTTTTAATTGACGTTGCAAAG |
| pLGB13-dBF9343_3444-3f |  | ACGTCAATTAAAAAACATGATTGTTTAGCAGTACAACAGTAATAAGATAAACGGTG |
| pLGB13-dBF9343_3444-3r |  | GAAGATAGGCAATTAGTCGACTTGGATAGGAACCCTTTTATTTTTTGCC |
